# RNA polymerase-mediated regulation of intrinsic antibiotic resistance and bacterial cell division

**DOI:** 10.1101/2025.07.20.665812

**Authors:** Vijay Soni, Junhao Zhu, Yesha Patel, John D. Helmann, Eric J. Rubin, Kyu Y. Rhee

## Abstract

Rifampin is a frontline antibiotic that inhibits the RNA polymerase of Mycobacterium tuberculosis (Mtb), the causative agent of tuberculosis (TB). Unlike most antibiotics, rifampin has an unusual ability to shorten the duration of treatment needed to cure TB that is not simply explained by its antimicrobial potency. We sought specific secondary effects of rifampin’s inhibition of Mtb RNA polymerase that may mediate this activity. We discovered that rifampin elicited a cell division arrest that was mediated through its inhibition of RNA polymerase. This arrest resulted in a downstream inhibition of the MtrAB two-component regulatory system, a mediator of intrinsic antibiotic resistance in Mtb. This inhibition is broadly conserved in other bacteria and represents a novel form of antimicrobial activity, termed adjunctive sensitization, that can mediate synergy and may contribute to rifampin’s unusual treatment shortening activity.

## INTRODUCTION

Rifampin (RIF) is an essential component of all current frontline treatments for TB, the leading cause of deaths not only due to an infectious but also curable, disease(*1*). This essentiality is a unique product of both its potent *in vitro* activity against *Mycobacterium tuberculosis* (Mtb)(*2*), the causative agent of TB, and its clinical ability to shorten the duration of treatment needed to achieve a durable, relapse-free cure(*3, 4*). However, resistance to rifampin, a defining feature of multi- (MDR), extensively- (XDR), and totally-drug (TDR) resistant TB, is on the rise, prompting a need for therapeutically equivalent alternatives(*1*).

Biochemical, structural, and genetic evidence has demonstrated that rifampin specifically binds the beta-subunit of bacterial RNA polymerase (RNAP) and blocks the exit tunnel of nascent transcripts to prevent their extension beyond a trinucleotide length and that this binding is required for antimicrobial activity(*5*) (Fig 1A). Knowledge of whether this antimicrobial activity is mediated or accompanied by specific secondary physiologic effects or is a non-specific consequence of inhibiting transcription initiation as a whole however remains unaddressed. Studies by Mitchison, for example, specifically showed evidence of continued incorporation of ^3^H uridine and recoverable Mtb mRNA following prolonged exposure to bactericidal concentrations of rifampin *in vitro*(*6*) while phenotypic studies revealed specific synergies between rifampin and β-lactam, but not protein synthesis, antibiotics(*7*). Mutations in RNAP conferring resistance to rifampin have been similarly reported to lead to changes in cell wall structure and bacterial morphology (Table S1).

**Figure 1:**
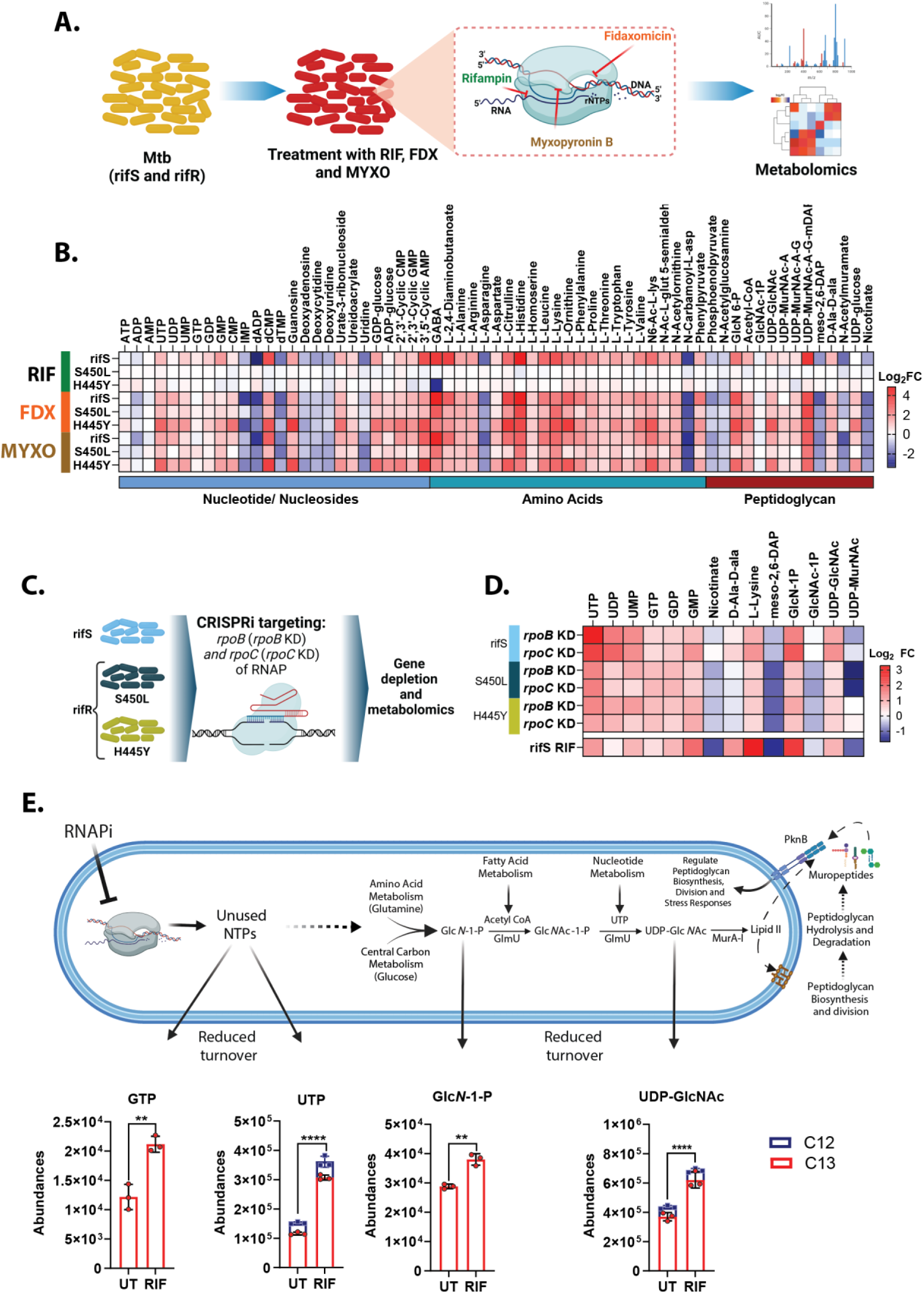
Inhibition of RNAP affects peptidoglycan metabolism of Mtb: **(A)** Experimental scheme used to define Mtb’s pre-lethal response to RNAP inhibitors (RNAPi). The inset shows the domain structure of RNAP highlighting the binding site of different RNAPi. Rifampin (RIF) binds at the exit channel. Fidaxomicin (FDX) binds at the base of the RNAP clamp. Myxopyronin B (MYXO) binds at the switch region of RNAP. **(B)** Heatmap showing statistically significant log_2_ fold changes (FC) in metabolite levels following a prelethal 10x MIC exposure to RNAP inhibitors both in rifS and rifR (S450L and H445Y) strains for 24 hrs. **(C)** Schematic of CRISPRi-mediated genetic inhibition of RNAP. sgRNA targeting *rpoB*, *rpoC*, and non-targeting (negative control) were electroporated into Mtb-Erdman (rifS and rifR) and gene knockdown phenotype was screened by growing mutants with and without anhydrotetracycline (ATc). Samples were collected for metabolomic profiling. **(D)** Heatmap highlighting log_2_ FC of metabolites upon CRISPRi-mediated silencing of RNAP subunits (RpoB and RpoC). Partial depletion of *rpoB and rpoC* was achieved using anhydrotetracycline (ATc) for 4 days, resulting in similar metabolic signatures as observed upon RNAP inhibitor administration (S2, and S3). Lower row displays matching metabolic changes of rifS-Mtb strain after RIF treatment. All results are representative of biological triplicates and two independent experiments. **(E)** Diagrammatic illustration showing the impact of RNA polymerase inhibition on peptidoglycan precursor and ribonucleotide pools in Mtb (top panel). Impact of rifampin (10x MIC dose; 0.5 µg/ml for 24 hrs) on pool sizes of GTP, UTP, Glc*N*-1P, and UDP-Glc*N*Ac in Mtb that had been prelabeled with ^13^C and transferred to ^12^C upon rifampin treatment. *p-*values were calculated using non-parametric t-test (for Glc*N*-1-P and GTP) or two-way ANOVA mixed model (for UTP and UDP-Glc*N*Ac). \*\*\*\**p*<0.0001, and **p<0.01. Error bars denote s.e.m.

Here, we provide evidence that rifampin-mediated inhibition of RNA polymerase in Mtb elicits specific physiologic changes in cell wall metabolism and division that contribute to its antimycobacterial activity. We further show that these effects are phylogenetically conserved in other bacterial species. These results thus reveal a previously unrecognized, but evolutionarily conserved, functional interaction between transcription and cell division that expands our understanding of the antimicrobial mode-of-action of a well validated molecular target.

## RESULTS

### Metabolic impact of RNAP inhibition

We previously demonstrated that, despite its bulk effect on RNA synthesis, rifampin elicited a specific set of metabolic changes that included a combination of some that were shared by other antibiotics and others that were specific to rifampin(*8*). We hypothesized that those specifically associated with the antimycobacterial activity of rifampin might serve as a biochemical window into its treatment shortening activity. We therefore compared the metabolic profiles of isogenic rifampin-sensitive (rifS) and -resistant (rifR) Mtb strains during the pre-lethal phase of exposure to rifampin (Fig 1A, S1A). Pathway enrichment analysis of such activity-specific metabolites revealed a statistically significant over-representation of metabolites associated with amino acids, nucleotides, peptidoglycan (PG) biosynthesis, and the tricarboxylic acid (TCA) cycle (Fig 1B, S1B, S1C). We validated the functional association of these changes with the inhibition of RNAP by profiling the metabolic responses of rifS and rifR strains of Mtb to two additional structurally and mechanistically distinct inhibitors of its RNAP, fidaxomicin (FDX) and myxopyronin B (MYXO), at similar levels of antimycobacterial activity (Fig 1B, S1D), as well as following partial transcriptional silencing of the β (*rpoB*) and β’ (*rpoC*) subunits of RNAP (Fig 1C, 1D, S2, S3). We further showed that these changes were not carbon source specific (S4A) and observed in both laboratory-adapted and clinical isolates (S4B). Metabolic tracing studies of Mtb following treatment with rifampin (S5) more specifically revealed accumulations of UDP-Glc*N*Ac, GTP, ATP, and UTP indicative of an increase in PG turnover and arrest of de novo PG biosynthesis (Fig 1E).

### Transcriptomic impact of RNAP inhibition

Seeking further evidence of a specific physiologic impact of rifampin, we analyzed the transcriptomes of rifS and rifR Mtb during the same pre-lethal phase of exposure to rifampin (Fig 2A, S6A). Despite its expected bulk inhibition of de novo transcription, treatment with rifampin elicited both increases and decreases in transcript abundance (Fig 2A). We next overlaid the transcriptional response of Mtb to rifampin with that of FDX, a structurally and mechanistically distinct inhibitor of RNAP to define a biologically more specific transcriptional signature of RNAP inhibition. Using an absolute cutoff (log_2_ fold>1 and *p_corr_* <0.05), we identified an activity-specific transcriptional signature that consisted in an accumulation of 537 and depletion of 344 transcripts (Fig 2A, S6B). Functional enrichment analysis revealed a depletion of genes associated with de novo PG biosynthesis (S6C), consistent with Mtb’s observed metabolic response. Further analysis of this signature revealed an enrichment of genes encoding and belonging to the *mtrAB* regulon that was not observed in rifR strains exposed to rifampin (Fig 2B, 2C). The *mtrAB* regulon consists of a two-component sensor kinase and cognate DNA-binding response regulator that was previously shown to regulate peptidoglycan remodeling enzymes required for cell growth and division and reported to mediate both drug tolerance and intrinsic drug resistance (Fig 2D)(*9–13*). Targeted qPCR analysis of *mtrAB* and a representative subset of its regulon demonstrated that this repression was specific to RNAP-targeting antibiotics (Fig 2E, S7) and carbon source independent (S8). This repression was additionally observed in Mtb whose replication had been slowed by hypoxia, acidic pH, or nutrient (PBS) starvation (S9).

**Figure 2:**
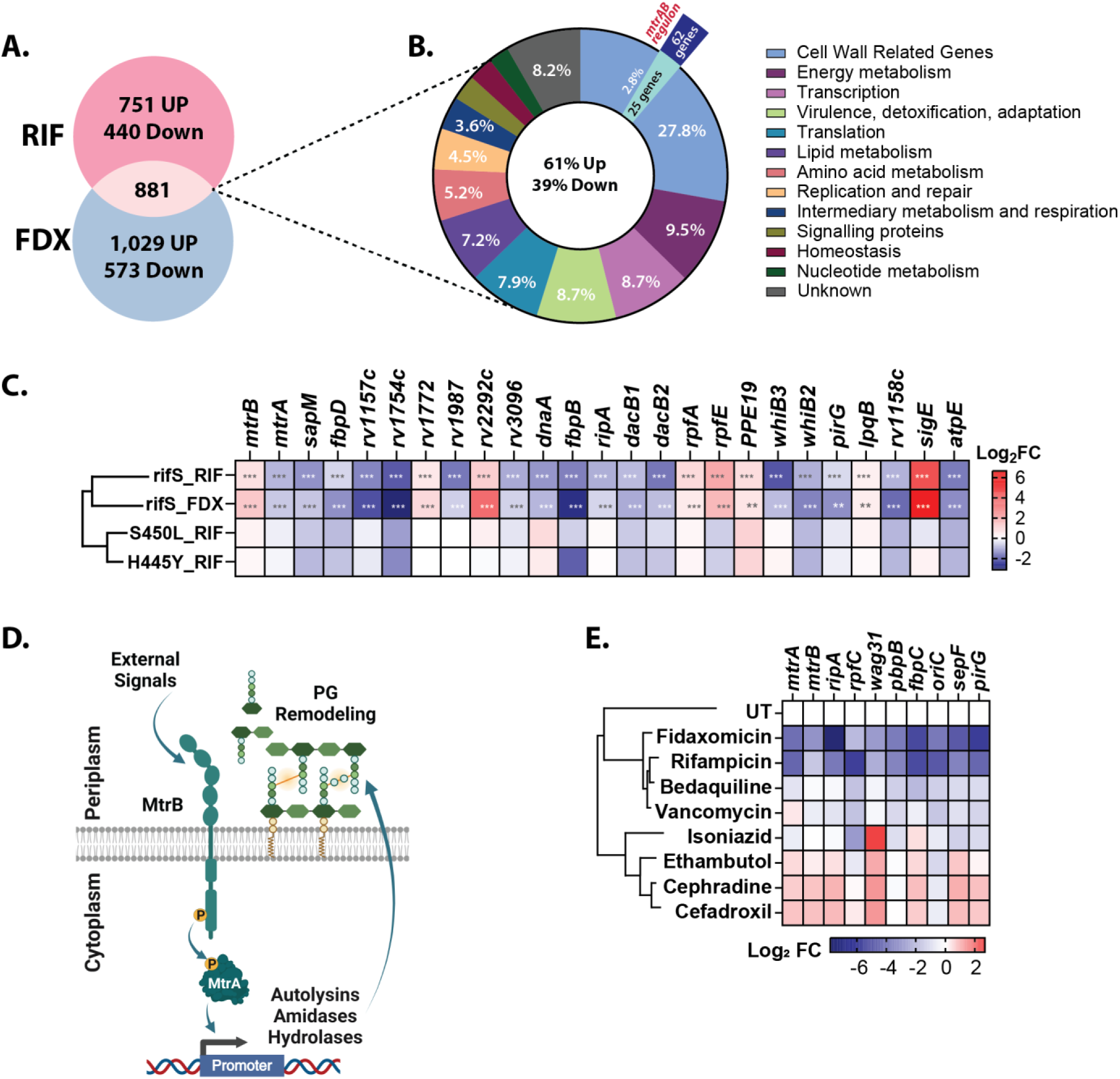
Transcriptomic profiles of Mtb RNAP inhibition by rifampin and fidaxomycin: **(A)** Venn diagram showing the differentially expressed and overlapping genes (in overlay) upon exposure of 10x MIC dose of rifampin and fidaxomicin to rifS strain (Mtb-H37Rv) for 24 hrs. Cutoff: log_2_ fold changes >1 and < -1; *p_adj_* value <0.05. **(B)** Donut graph shows the functional enrichment analysis of significant overlapping genes with known functions. Almost 27.8% of genes belong to the cell wall-related mechanism and 2.8% of total genes belong to *mtrAB* regulon (see overhang) **(C)** Heatmap displaying log_2_ fold changes of representative members of the *mtrAB* regulon upon inhibition of RNAP both in rifS and rifR strains. Stars show the range of *p_adj_* values for the most significant differential expressions calculated using the Benjamini-Hochberg method. *p* values: **< 0.05; ***<0.0005. **(D)** Depiction of the *mtrAB* regulon and its function in bacteria. Autophosphorylation or external stimuli (unknown) activates the MtrB sensor kinase which results in the downstream phosphorylation-mediated activation of its cognate response regulator MtrA and expression of almost 62 genes participating in PG remodeling, maintenance, cell shape, elongation, division, and intrinsic antimicrobial resistance (AMR). **(E)** Real-time qPCR results showing log_2_ fold changes of some key genes from *mtrAB* regulon after treatment of different RNAP inhibitors and antibiotics targeting cell wall or respiration. Centroid linkage hierarchical cluster analysis was done using Gene Cluster 3.0 software. All results are representative of biological triplicates and two independent experiments.

### Effect of RNAP on divisome activity

Given the foregoing data, we sought to probe the functional relationship between RNAP activity and cell wall metabolism in replicating Mtb. Prior studies reported a robust but mechanistically unexplained phenotypic synergy between rifampin and cephalosporin antibiotics against Mtb(*7*). We found that this synergy is also extended to a panel of cell wall-targeting a β-lactam antibiotics (cephradine, cefadroxil, meropenem, and faropenem) that selectively inhibit FtsI (PbpB or Rv2163c)(*14–20*), the transpeptidase required for septal peptidoglycan biosynthesis during cell division(*21, 22*), but not inhibitors of other peptidoglycan biosynthesis, including vancomycin and cycloserine (Fig 3A).

**Figure 3:**
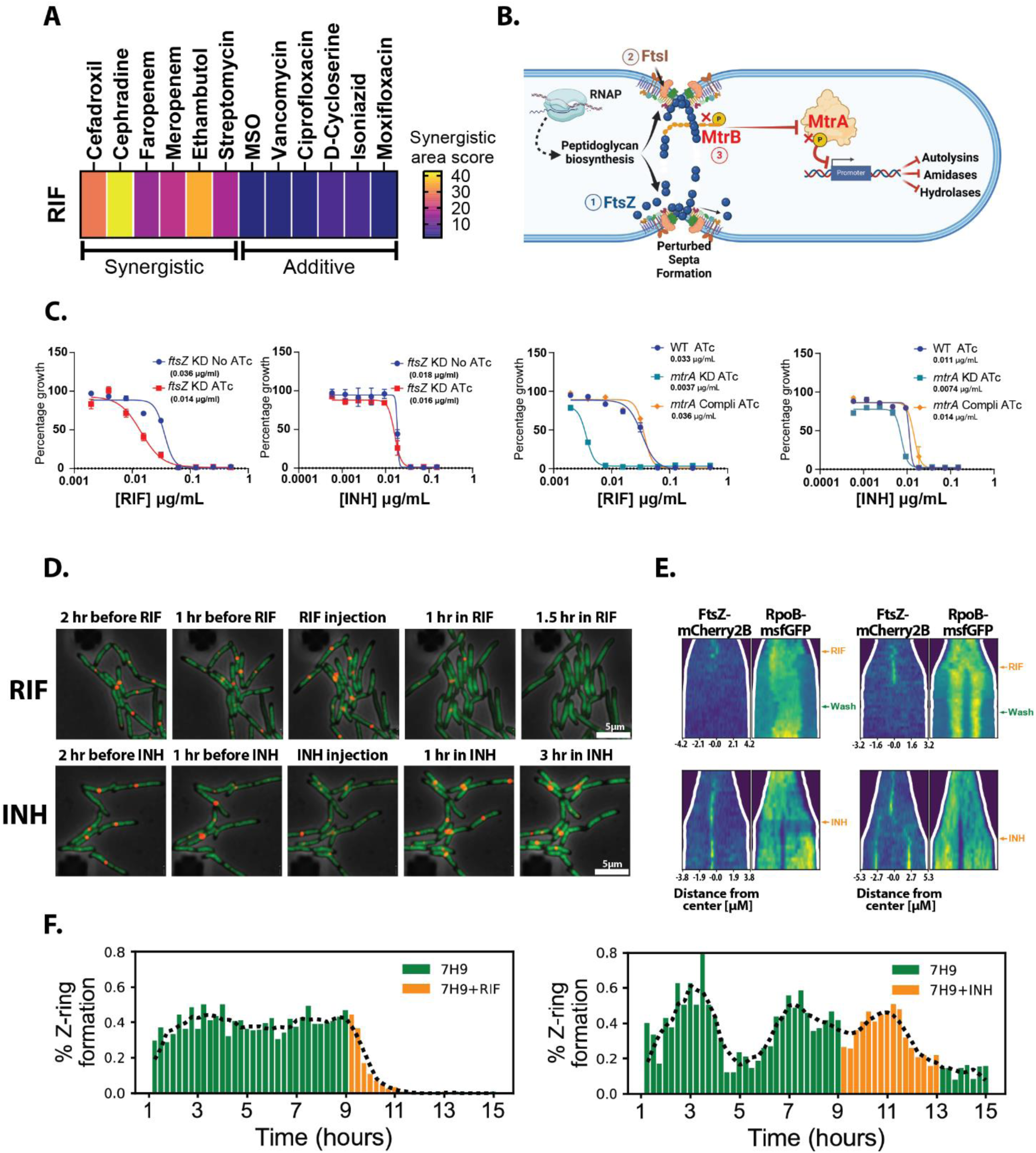
Impact of RNAP inhibition on divisome assembly: **(A)** Heatmap showing the most synergistic area score of rifampin combined with antibiotics of differing modes of action. Most synergistic area scores and Zero Interaction Potency (ZIP) scores were measured using SynergyFinder (https://synergyfinder.fimm.fi/). Scale: less than -10: antagonistic, from -10 to 10: additive, larger than 10: synergistic. All experiments were done in biological triplicates. (Table S3). **(B)** Graphic showing the impact of localization of the divisome components FtsZ (*1*) and FtsI (*2*) on the activation of MtrBA (*3*) system and their role in peptidoglycan biosynthesis and remodeling. Numbers (1,2, and 3) show the sequence of molecular events. **(C)** Dose-response graphs showing the minimum inhibitor concentrations of rifampin (RIF) and isoniazid (INH) upon partial depletion of *ftsZ* and *mtrA* in Mtb-Erdman. Gene knockdown was achieved using anhydrotetracycline (ATc) regulated CRISPRi constructs expressing respective sgRNA. **(D-F)** Subcellular localization dynamics of FtsZ and RpoB following RIF and INH treatment. Dynamics of FtsZ and RpoB were tracked in *M. smegmatis* mc^2^155 cells expressing fluorescent fusions of RpoB (rpoB-msfGFP) and FtsZ (ftsZ-mCherry2B) (S13). Bacteria were grown for 9 hrs followed by 4 hrs of drug treatment (20µg/ml RIF and 25µg/ml INH) and 2 hrs of recovery. Spatial-temporal expression dynamics of the fluorescent proteins (in yellow color) were measured using Fiji and customized Python scripts. (D) Representative microscopy images illustrating Z-ring dynamics upon RIF or INH treatment. (E) Single bacteria kymographs highlighting differences in Z-ring formation between RIF and INH treated cells. The upper left panel shows that RIF inhibits FtsZ localization, whereas the right panel reveals that RIF can also halt FtsZ recruitment even after cell division has initiated. (F) Bar charts showing the percentage of cells with detectable FtsZ foci at each time point. Scale bar: 5µm. Results represent biological triplicates across two independent experiments, except for time-lapse imaging, which represents two independent experiments with biological duplicates.

Previous work had demonstrated that MtrB, the sensor kinase, localized to the cell septum via interaction with divisome components (FtsI and FtsZ) and that this interaction was required for activation of *mtrA* and its regulon(*23, 24*). We, therefore, sought to test if the impact of rifampin on *mtrAB* regulon activity might be a downstream consequence of impaired septal Z-ring formation (Fig 3B). To do so, we first tested the effect of inhibiting Z-ring formation on rifampin susceptibility by transcriptionally silencing the expression of *ftsZ* to levels that only mildly slowed but did not arrest growth (S10). This revealed a selective sensitization to rifampin, but not isoniazid, upon *ftsZ* silencing (Fig 3C, S10). Similar effects were observed upon *mtrA* knockdown (Fig 3C, S11). We further found that treatment of Mtb with FtsI inhibitors resulted in increased levels of UDP-GlcNAc (S12).

Seeking more direct evidence of the effect of rifampin on septal Z-ring assembly, we conducted single-cell time-lapse microscopy of reporter strains that either expressed fluorescent fusions of *ftsZ* and *rpoB* and were exposed to defined pulses of rifampin (S13). Owing to resource limitations and a high degree of conservation of both FtsZ and RpoB in Mtb and *M. smegmatis* (89.61% amino acid identity/94% similarity and 99% amino acid identity/95% similarity, respectively), we conducted these experiments in *M. smegmatis*(*25, 26*) (S14 and S15). These studies demonstrated that rifampin selectively inhibited Z-ring formation in cells that had recently or not yet initiated cell division while allowing the remainder to complete the process of chromosomal segregation and cytokinesis (Fig 3D and E, upper panels). In contrast, treatment with isoniazid revealed no such effect on Z-ring formation (Fig 3D and E, lower panels), though both isoniazid and rifampin slowed cell elongation (Fig 3E, 3F). We further quantified the fraction of cells with clear FtsZ-mCherry2B foci at different stages of cell division and found that rifampin exposure almost completely collapsed nascent Z-ring formation within 2 hours of exposure, whereas isoniazid permitted continued Z-ring assembly (Fig. 3F).

### Conservation of RNAP-MtrAB interaction

The biological centrality and evolutionary conservation of both RpoB and FtsZ across bacteria taxa further led us to wonder if the foregoing effects of RNAP inhibition might also be conserved in other bacterial species (S15). Focusing specifically on bacteria genera annotated to encode orthologs (WalKR/YycFG/VicKR) of the MtrAB two-component regulatory system (Fig 4A and B), we characterized the molecular and phenotypic effects of rifampin against *Bacillus subtilis* (BS) and *Staphylococcus aureus* (SA). Both organisms encode orthologs of *rpoB* and *ftsZ* with greater than 55% amino acid identity and 75% similarity (S15) and MtrAB orthologs exhibiting greater than 45% identity and 66% similarity (S16) with similar roles in cell wall remodeling, division, and antibiotic resistance(*27–31*) (Table S2). Moreover, exposure to rifampin, fidaxomicin and myxopyronin B in both organisms demonstrated evidence of increased levels of UDP-Glc*N*Ac (Figure 4C, S17, S18), repression of their respective *mtrAB* orthologs (Figure 4D), and phenotypic synergy with cephalosporin and carbapenem antibiotics (Figure 4E).

**Figure 4:**
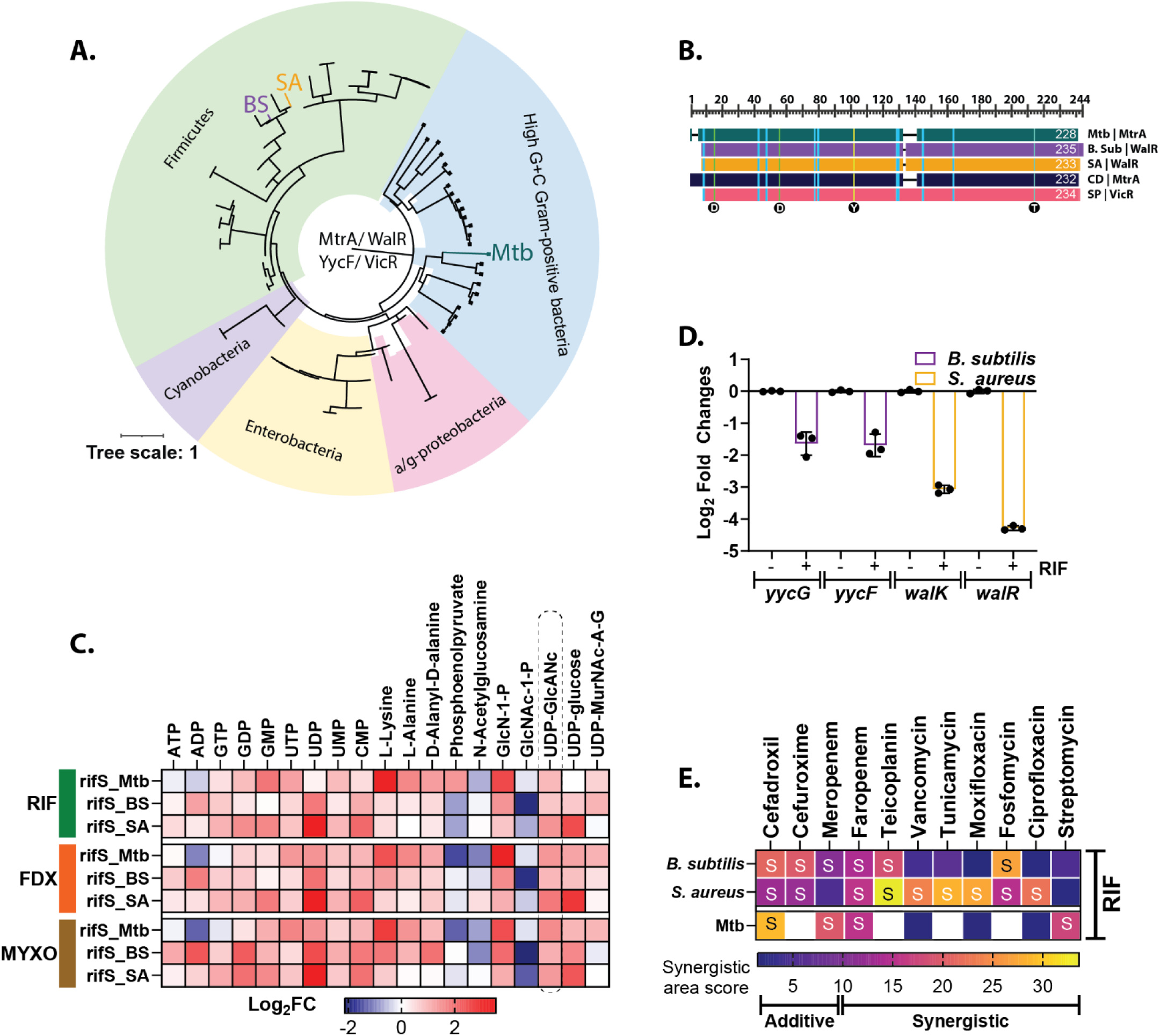
Phylogenetic conservation of the RNAP-cell wall-division relationship: **(A)** A phylogenetic tree of MtrA orthologs in different bacterial species was created using https://itol.embl.de/. **(B)** Graphic presenting the conserved two-domain structure of MtrA and its orthologs. Conserved aspartic acids on the N-terminal domain and tyrosine on the C-terminal domain are targets of the sensor kinase (MtrB or WalK or YycF), while threonine on the C-terminal domain is primarily targeted by a serine-threonine protein kinase (PknB or PrkC). **(C)** Heatmap representing log_2_ FC of key metabolites in *B. subtilis* and *S. aureus* upon treatment with RIF, FDX, and MYXO for 20 min (S17). Respective metabolites from rifS-Mtb treated with RIF are also presented for as reference. **(D)** Real-time qPCR estimation showing the levels of two-component systems upon RIF treatment for *B. subtilis* and *S. aureus walKR*. A significant depletion is consistent with changes in the corresponding MtrAB regulon of Mtb. **(E)** Distribution of most synergistic area score showing the extent of synergy of RIF with different antibiotics both in *B. subtilis* and *S. aureus.* Lower segment is showing the overlapping antibiotic synergy score from rifS-Mtb strain. White cells indicate an absence of corresponding antibiotics. All results are representative of biological triplicates and two independent experiments.

## DISCUSSION

Owing to the biological centrality of transcription and well-defined molecular mechanism of biochemical inhibition, studies of rifampin have historically focused on its quantitative impact on rates of transcription initiation as a whole, rather than the specific physiologic consequences of that inhibition(*5*). Our studies of both rifamycin and non-rifamycin inhibitors of RNA polymerase against rifampin-sensitive and -resistant strains now shed new light on the latter. Despite its genome-scale impact, we discovered that inhibition of transcription initiation, or genetic silencing of the β- or β’ subunits of RNA polymerase, elicited physiologically specific and reversible metabolic changes that tracked with its antimycobacterial activity, and were associated with both transcriptional and phenotypic inhibition of the *mtrAB* two-component regulatory system (Fig 5). This inhibition effectively sensitizes Mtb (not completely killed by inhibition of transcription) to a number of secondary host- and drug-imposed stresses, and, in doing so, reveals a previously unrecognized form of antimicrobial impact, adjunctive sensitization, that expands the scope of rifampin’s antimicrobial mode-of-action.

**Figure 5:**
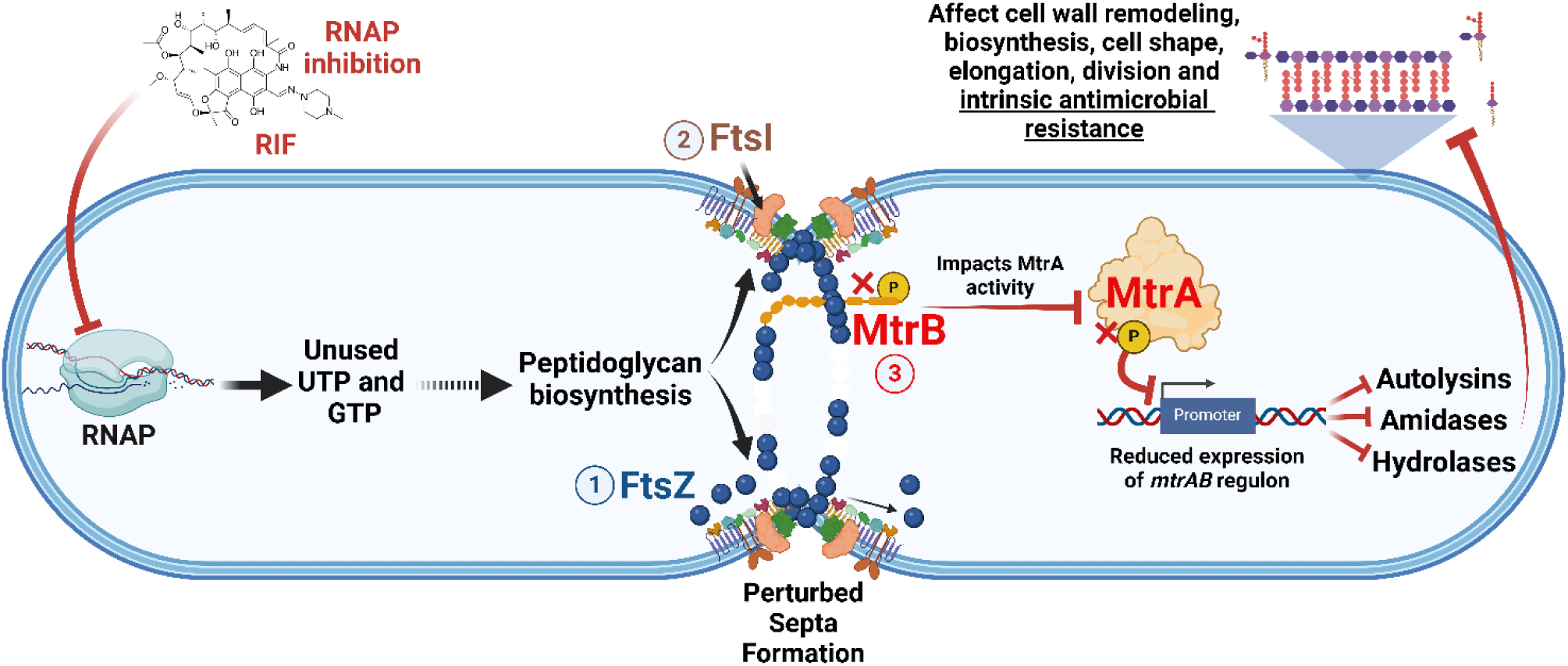
Generalized model of RNAP mediated regulation of cell wall biosynthesis and bacterial division: Under normal growth conditions, bacteria maintain a state of metabolic homeostasis that provides a continuous supply of nucleotides to sustain transcription and cell wall biosynthesis. Perturbations of transcription (either antibiotic or genetic) result in diminished turnover of nucleotides (UTP and GTP) that may affect de novo peptidoglycan biosynthesis and septa formation. The resulting imbalance leads to septum delocalization, consequently suppressing the regulon of the bacterial two component system (MtrAB/YycFG/WalKR or VicKR) and disrupting cell wall biosynthesis, elongation, bacterial division, and intrinsic antimicrobial resistance.

Among currently approved TB drugs, rifampin is distinguished by its ability to shorten the duration of treatment needed to achieve a durable, relapse-free cure(*32*), an activity not simply explained by standard measures of its *in vitro* antimycobacterial potency(*33, 34*). This activity has instead been ascribed to a combination of its ability to penetrate caseous or necrotic lesions and kill the non- or slowly replicating Mtb subpopulations therein(*35–37*). However, the mechanistic basis of this activity remains incompletely defined. That rifampin’s impact on *mtrAB* was both carbon source independent and extended to non- or slowly replicating Mtb populations highlights a novel mechanistic effect of high potential therapeutic importance. Treatment shortening aside, the potential for this activity to inform the development of rational mechanism-based drug combinations warrants further study.

Translational potential notwithstanding, this work extends our understanding of the fundamental biology of RNA polymerase. Previous studies of the *mtrAB* and *walKR* systems had demonstrated its interaction with and regulation of genes involved in cell division and cell wall metabolism, many of which include enzymes involved in peptidoglycan remodeling(*10–12, 29, 38, 39*). In Mtb and other bacteria, MtrAB or WalKR activity has been shown to depend on septal localization and therefore be functionally associated with Z-ring formation and cell division(*23, 24, 38–41*). As discussed previously, mutations conferring resistance to RIF (in the RpoB subunit of RNAP) in other microbes have been reported to changes in cell wall structure(*42*), bacterial morphology(*42–45*), virulence(*46*) as well as resistance to various drugs targeting peptidoglycan biogenesis (e.g., vancomycin and daptomycin)(*47–50*) while resistance to some β-lactams and cephalosporins has conversely been reported to be associated with mutations in the *rpoB* and *rpoC* subunits of RNAP(*7, 42, 43, 51–54*) (*55, 56*) (Table S1). Our work now extends this biology further to include an even more upstream and evolutionarily conserved role of RNA polymerase and its state of transcriptional competence as a potential checkpoint regulator of cell division.

Growing evidence has implicated a broader range of physiologic roles for RNA polymerase beyond its activity as a bulk enzymatic catalyst of RNA synthesis(*43, 51, 57*). Once focused on the primary drug-target interaction, modern drug development has expanded in scope to include studies of specific secondary or downstream consequences of the drug-target interaction. Such studies have helped increase knowledge of the mechanistic basis for drug activity, and in doing so, reveal additional new potential targets whose inhibition could mimic the activity of the drug itself and/or slow/prevent the emergence of resistance. Though less well recognized, such studies have also created a window into the normal physiologic functions of the targets they inhibit.

## Acknowledgments

We thank to TBRU and TB-alliance for providing various RNAP inhibitors in this study. We thank Jenny Zhaoying Xiang and Adrian Y Tan from the Genomics core of Weill Cornell Medicine for their constant support for RNA sequencing and genomics analysis. We acknowledge Dirk Schnappinger, and Jeremy Rock for providing CRISPRi strains of *ftsZ* and *mtrA*. We also thank Michael DeJesus, Nicholas Poulton, Shuqi Li, Dr. Cara Boutte, Kristin Burns-Huang, Allison Fay, Daniel Fitzgerald, David Alland, David Sherman, Murty Madiraju, and Dr. Dhandayuthapani S. for their help with different clinical, lab, and drug-resistant strains. Thanks to Jennifer Herrmann from Helmholtz Institute for Pharmaceutical Research Saarland, Germany for generously providing Myxopyronin B.

## Funding

K.Y.R.: National Institutes of Health grant: U19 AI162584 K.Y.R.: Bill & Melinda Gates Foundation: BMGF INV-004709 J.D.H.: National Institutes of Health grant: R35GM122461

## Author contributions

Conceptualization: V.S. and K.Y.R.

Methodology: V.S., K.Y.R., J.Z.

Investigation: V.S. and K.Y.R.

Visualization: V.S. and K.Y.R.

Funding acquisition: K.Y.R.

Project administration: K.Y.R.

Supervision: K.Y.R.

Writing – original draft: V.S. and K.Y.R.

Writing – review & editing: V.S., K.Y.R., Y.P, J.D.H., and E.J.R.

## Competing interests

The authors declare that they have no competing interests.

## Data and materials availability

Raw metabolomics data are available at Metabolomics Workbench under the project ID [PR002179](https://dev.metabolomicsworkbench.org:22222/data/DRCCMetadata.php?Mode=Study&DataMode=AllData&StudyID=ST003543&StudyType=MS&ResultType=1&Access=UxyM1248#DataTabs). Additionally, the raw transcriptomics data are deposited to the NCBI Short Read Archive with the BioProject ID [PRJNA1181268](https://www.ncbi.nlm.nih.gov/bioproject/1181268). All code used to generate cell kymographs and representative micrographs was deposited on GitHub and available at https://github.com/jzrolling/FtsZ_kymographs. Other data are available in the supplementary materials.

## Abbreviations

Tuberculosis: TB
*Mycobacterium tuberculosis*: Mtb
Multi-drug resistance: MDR
Extensive-drug resistance: XDR
Total-drug resistance: TDR
RNA Polymerase: RNAP
Rifampin: RIF
Rifampin-sensitive: rifS
Rifampin-resistant: rifR
UDP-GlcNAc: UG
Peptidoglycan: PG
Tricarboxylic acid: TCA

## Notes

### Competing Interest Statement

The authors have declared no competing interest.

