## Supplementary for "RNA polymerase-mediated regulation of intrinsic antibiotic resistance and bacterial cell division"

### MATERIAL AND METHODS

#### Chemicals

Different antibiotics used in this study are listed in Table S7.

#### Bacterial Strains

Bacterial strains used in this work are listed in Table S4, respectively. Mtb strains H37Rv, Erdman, CDC1551, SA161, Haiti-17874, Haiti-20142, Haiti-20389, rifampin-resistant strains (S450L and H445Y), and CRISPRi mutants (of *rpoB*, *rpoC*, *ftsZ,* and *mtrA*) were maintained and cultured in a biosafety level 3 facility at 37 °C using Difco$^{TM}$ Middlebrook 7H9 or 7H10 medium supplemented with 0.2% glycerol and 10% ADN containing 0.85 gm/lit NaCl, 2 gm/lit D-glucose, and 5 gm/lit of Fraction V BSA (Roche). Similarly, *M. smegmatis* mc$^2$155 wild-type and recombinant strains expressing fluorescent fusions of *ftsZ* or *rpoB* were cultured on 7H9 and 7H10 media as mentioned above. Both *B. subtilis* st 168 and *S. aureus* N315 were cultured on Difco$^{TM}$ LB broth medium at 37°C aerated on an orbital shaker at 200 rpm. *B. subtilis* rifampin resistance mutants were used from the previous study(*43*).

#### Bacterial Culture Conditions

Mtb metabolomics, isotope labeling, transcriptomics, qPCR, and stress experiments were done using bacteria-laden filters. For this, 1 ml of mid-log phase ($OD_{580}$ = 0.8–1) planktonic culture of each strain was passed through 0.22-µm nylon filters, retaining only bacteria. These filters were placed on Middlebrook 7H10 agar (BD) supplemented with 0.2% glycerol and ADN and allowed to grow Mtb for the next 5 days at 37 °C. To acclimatize, on day 6, the bacteria-laden filters were moved onto reservoirs filled with 7H9 medium (no tyloxapol) and incubated for 24 hrs at 37 °C. Next day, these bacteria were exposed to different antibiotics (at 1x, 5x, and 10x MICs concentrations) or ATc (100 ng/ml) or other stressors for another 24 to 48 hr or more (for labeling and time course) at 37 °C. Samples (bacteria and supernatants) were collected and processed by different methods. *B. subtilis* (both rifS and rifR strains) and *S. aureus* were cultured up to mid-log phase (0.5 $OD_{600}$) in LB broth at 37 °C using an orbital shaker. 10-30 ml of these cultures were exposed to different antibiotics for 20 min at 37 °C under static conditions. These cultures were then collected at 4,000 RPM and 4°C for 10 min. All experiments were conducted in triplicates across two or more independent experiments.

#### Generation of CRISPRi Knockdown

Different CRISPRi were cloned using previously published methods(*58*). PEBBLE (https://pebble.rockefeller.edu/tools/sgrna-design/) platform was used to design sgRNAs targeting the non-template strand of the target gene open reading frame. pLJR965 (Addgene #115163) plasmid backbone was digested with BsmB1-v2 (NEB R0739L) and ligated with sticky end overhangs of two annealed complementary oligonucleotides of sgRNAs using T4 ligase (NEB M0202M). Successful clones were confirmed by Sanger sequencing. Each CRISPRi plasmids were electroporated into Mtb strains. For electrocompetent bacteria, Mtb strains were cultured up to $OD_{580}$ = 0.8-1.0 and centrifuged at 4,000 RPM for 10 min at room temperature (RT). Bacterial pellets were then washed (4,000 RPM, 10 min, RT) three times with sterile 10%

glycerol and then resuspended in 10% glycerol to a final volume that was 10% of the original culture volume. Next, 100 µl of electrocompetent Mtb strains were mixed with 200 ng CRISPRi plasmid and transferred into a 2 mm electroporation cuvette (Bio-Rad 1652082). Bio-Rad Gene Pulsar X Cell electroporation system was used for the electroporation, with settings of 700 Ω, 2,500 V, and 25 µF. Transformed Mtb was recovered in 7H9 broth for 24 hrs at 37 °C followed by plating on 7H10 agar plates with kanamycin (50 µg/ml) to select positive transformants. Final Mtb strains with different gene knockdowns were grown with or without ATc (100 ng/ml) and confirmed using qPCR.

**MIC and Synergy**

To determine the 1x MIC dose for multiomics experiments, 1 ml Mtb culture of $OD_{580}$ = 0.8 was passed through 0.22-µm nylon filters and placed on reservoirs containing 7H9 media with different dilutions of antibiotics. Visible growth was assessed after 7 and 10 days. The lowest concentration of antibiotics that completely inhibited the bacterial growth was determined as 1x MIC dose. For antibacterial activity assays, all antibiotics were resuspended in DMSO or water, and serial dilutions were aliquoted using an HP D300e digital dispenser in a 96-well plate format. While, for the checkerboard assay, a matrix of two antibiotics concentration ranges were aliquoted across columns and rows of 96 well plates. The maximum concentration of DMSO was 1% of the final volume of culture and maintained at the same concentration in all wells. Different Mtb strains were grown up to the mid-log phase ($OD_{580}$ = 0.8) while CRISPRi knockdown strains were growth synchronized and pre-depleted using ATc (100 ng/ml) for 5 days. *B. subtilis* and *S. sureus* were grown up to $OD_{600}$ = 0.5. All bacterial cultures were diluted to a starting $OD_{580}$ or $OD_{600}$ of 0.02 in 7H9 media or LB broth and 200 µl bacterial suspension was dispensed in technical triplicates in all wells containing fresh ATc and/or antibiotics. Plates were incubated statically at 37 °C with 5% $CO_2$. Mtb plates were assessed on days 10 and 14, and $OD_{580}$ was measured using Tecan Spark plate reader. While for non-Mtb bacteria $OD_{600}$ was measured 18 and 24 hrs post plating. Percentage growth was calculated with respect to DMSO control for each strain. $IC_{50}$ were computed using nonlinear fit curves in GraphPad Prism 10 software. All MIC curves represent mean ± s.e.m. of all triplicates and are representative of a minimum of two independent experiments. Synergy was determined using Zero Interaction Potency (ZIP) scores and most synergistic area scores calculated by SynergyFinder software (https://synergyfinder.fimm.fi/)(*59, 60*). A ZIP score of more than 10 represents synergy between the two antibiotics.

**Metabolomics**

Metabolic profiling and analysis were performed as published before(*43, 61, 62*). In brief, bacteria laden Mtb filters were exposed to 1x, 5x, and 10x MIC doses of antibiotics for 24 hrs. While *B. subtilis* and *S. aureus* were grown planktonically and 30 ml of 0.5 $OD_{600}$ cultures were treated with antibiotics for 20 min. Samples were quenched in a 1ml precooled (-40 °C) mixture of acetonitrile, methanol, and water (40%:40%:20%). Metabolite extractions were done by bead beating using 0.1 mm Zirconia beads and Precellys homogenizer (Bertin Technologies) for 3 min (6,000 RPM, 3 rounds) with constant colling at or below 1°C. Lysates were cleared by centrifuging (13,000 rpm, 10 min, 4 °C) and sterilized by passing through Spin-X® Centrifuge tube filters (0.22 µm; Sigma–Aldrich).

To maximize biochemical coverage of different metabolite classes, two different liquid chromatography methods were employed (a) formic acid (FA)-based chromatography and (b) hydrophilic interaction liquid chromatography (HILIC). For FA method, 2 µl of extracted and cleaned samples were resolved on a Diamond Hydride Type C Column (Cogent; Catalog Number: 70000-15P-2) using a 1260 Infinity II high-performance liquid chromatography (HPLC) system (Agilent) coupled with an Agilent Accurate-Mass 6230 TOF-Mass Spectrometer (MS) operating both in negative and positive ionization modes. For a broad range of biochemical separation two liquid phases, (a) solvent A (water mixed with 0.2% FA), and (b) solvent B (acetonitrile mixed with 0.2% FA were used at different gradients with a flow rate of 0.4 ml/ min: 85% B (0 to 2 min); 80% B (3 to 5 min); 75% B (6 to 7 min); 70% B (8 to 9 min); 50% B (10 to 11 min); 20% B (11 to 14 min); 5% B (14 to 24 min) and 10 min of re-equilibration period using 85% B. For the HILIC method, 3 µl of samples were resolved on Agilent 1290 Infinity HPLC system using an Agilent InfinityLab Poroshell 120 HILIC-Z (Agilent 683775-924), 2.7 µm particle size, 2.1 × 150 mm column heated to 50 °C. Two solvent phases were used for the HILIC method (a) solvent A (100% water mixed with 10 mm ammonium acetate, 5 mM InfinityLab Deactivator Additive, pH 9 was set using $NH_4OH$), and (b) solvent B (85% acetonitrile mixed in water and 10 mM ammonium acetate, 5 mM InfinityLab Deactivator Additive, and pH 9 was set using $NH_4OH$). At a flow rate of 0.250 ml/min, both solvents were used with the following gradients: 96% B for 0 to 2 min; 88% B for 5.5 to 8.5 min; 86% B for 9 to 14 min; 82% B for 17 min, 65% B for 23 to 24 min; 96% B for 24.5 to 26 min; and the end-run at 96% B for 10 min. Data were collected on an Agilent 6230 Time of Flight (TOF) mass spectrometer with Agilent Jet Stream electrospray ionization (ESI) source (operated at 3500 V Cap, 0V nozzle voltage, gas temperature: 325 °C, drying gas: 8 L $min^{-1}$, nebulizer: 45 psig, sheath gas temp: 400 °C, sheath gas: 12 L $min^{-1}$, Vcap: 4000 V, and Fragmentor: 125 V) in extended dynamic ranges and either in both modes (for FA) or negative mode (for HILIC). Data collection was in centroid mode for m/z values ranging from 50 to 1700 Dalton. Metabolites were identified based on exact mass-retention time identifiers for peaks displaying the anticipated distribution of associated isotopes. Peak abundances of all metabolites were analyzed and estimated using Profinder 8.0 (Agilent) and Qualitative Analysis B.06.00 (Agilent) software and the KEGG metabolite database as a reference. To ensure the accuracy of identified peaks, standard metabolites were included in each run, and peak alignments were performed. Absolute and relative counts (with respect to untreated or wild-type controls) were calculated and heatmaps were plotted using Cluster 3.0, Java TreeView, and GraphPad Prism software. Each dose was analyzed in triplicates and the results are representative of two or more independent experiments.

### Metabolic Labelling

For selective $^{13}C$ labeling of Mtb metabolome, Mtb was first cultured on modified 7H9 containing [$^{13}C$] glucose, [$^{13}C$] glycerol, and [$^{13}C$] glutamate (Cambridge Isotope Laboratories) for 3-4 division cycles followed by filter preparations as discussed above. All detected metabolites were more than 95% [$^{13}C$] labeled (S5) These fully labeled bacteria-laden filters were transferred onto reservoirs containing fresh 7H9 media with unlabeled glucose, glycerol, and glutamate. Samples were collected at different time points and [$^{13}C$] labeling pattern was estimated using LC-MS as described in the metabolomics section. Percentage replacements of [$^{13}C$] labeled metabolites with [$^{12}C$] metabolites were calculated using the isotopologue extraction feature of Profinder software (Agilent Technologies).

**Stress Experiments**

Actively growing Mtb culture (50 ml of each replicate of $OD_{580}$ = 0.8 to 1.0) was washed twice in PBS-0.02% tyloxapol (PBST) and subjected to various stresses (S9). For PBS-based nutrient starvation, washed pallets were resuspended in PBST and incubated for 7 days at 37°C. On day 8$^{th}$ starved Mtb was exposed to different antibiotics. For pH stress, washed Mtb pellets were resuspended in 7H9 media with pH 5.5 while for hypoxia Mtb cultures were resuspended in 7H9 media and incubated at 0.1% $O_2$ at 37°C. For nonreplicating (NR) stress, washed Mtb pellets were resuspended in nonreplicating (NR) media. NR media consist of modified Sauton's medium as base (in 1 lit 0.5 gm $KH_2PO_4$, 0.05 gm ferric ammonium citrate, 0.5 gm $MgSO_4$, 0.085% NaCl, 0.5% bovine serum albumin (BSA), and 0.02% tyloxapol) at pH 5.0 along with 0.5 mM $NaNO_2$, 0.05% butyrate as sole carbon source and incubated at 1% $O_2$ and 5% $CO_2$ at 37°C. Bacteria were preadapted for 24 hours in hypoxia, pH, and NR stress. After achieving different stress states, Mtb cultures were exposed to 1x and 50x MIC doses of rifampin (RIF) and isoniazid for two days and collected by centrifugation (4,000 RPM, 10 min, RT) and processed for RNA isolation and qPCR analysis.

**Transcriptomic Experiments and Analysis:**

*Total RNA Isolation:* Mtb-laden filters or planktonic cultures (20 ml, $OD_{580}$ = 0.6 to 1.0) were collected in 1 ml GTC buffer (5 M guanidinium thiocyanate, 25 mM trisodium citrate dihydrate, 0.5% Na- N-lauroylsarcosine, and 0.1 M β-mercaptoethanol), followed by centrifugation (4,000 RPM, 10 min, RT) (S6A). Pellets were resuspended in 1 ml TRIzol reagent (Invitrogen, Thermo Fisher 15596026) and lysed using 0.1 mm zirconium beads. Lysed samples were cleared by centrifugation (13,000 RMP, 10 min, 4 °C), and 200 µl of chloroform was added and stored at -80 °C until the next step. For total RNA extraction samples were thawed and centrifuged to separate organic and aqueous phases. Next 500 µl 99.90% isopropanol (Fisher Chemical A4611) was added to precipitate total nucleic acid and the pellet was washed with 99.45% ethyl alcohol (Sigma, E7023). Genomic DNA was digested using TURBO DNase (Invitrogen, AM1907). DNase was removed by an RNA cleanup kit (Zymo Research R1017) and concentrated RNA was estimated using nanodrop.

*RNA-Seq (rRNA depletion method for TB):* Following RNA isolation, total RNA integrity was checked using a 2100 Bioanalyzer (Agilent Technologies, Santa Clara, CA). RNA concentrations were measured using the NanoDrop system (Thermo Fisher Scientific, Inc., Waltham, MA). RNA sample library and RNA-seq were performed by the Genomics Core Laboratory at Weill Cornell Medicine. Firstly, rRNA was removed from Total RNA using the Qiagen QiaSeq Fast Select kit. Following purification, the Illumina TruSeq Stranded Total RNA Sample Library Preparation kit was used (Illumina, San Diego, CA), according to the manufacturer's instructions. Messenger RNA was fragmented into small pieces using divalent cations under elevated temperature. The cleaved RNA fragments were copied into first-strand cDNA using reverse transcriptase and random primers. Then strand of mRNA was removed by RNaseH and a replacement strand was synthesized, incorporating dUTP in place of dTTP to generate ds cDNA. The incorporation of dUTP quenches the second strand during amplification. The cDNA fragments then went through an end repair process, the addition of a single ‘A’ base, and then ligation of the adapters. The products were then purified and enriched with PCR to create the final cDNA library. The normalized cDNA libraries were pooled and sequenced on Illumina NovaSeq6000 sequencer with pair-end 100 cycles. The raw sequencing reads in BCL format are processed through bcl2fastq 2.19 (Illumina) for FASTQ conversion and demultiplexing.

*Analysis of transcriptomics data:* The sequencing libraries were sequenced with paired-end 50 bps on the NovaSeq6000 sequencer. The raw sequencing reads in BCL formats were processed through bcl2fastq 2.20 (Illumina) for FASTQ conversion and demultiplexing. After trimming the adaptors with cutadapt (version1.18) ( https://cutadapt.readthedocs.io/en/v1.18/), RNA reads were aligned and mapped to the Mtb genome (H37Rv or Mtb-Erdman) by Bowtie 2 version 2.2.8 (http://bowtie-bio.sourceforge.net/bowtie2/index.shtml)(*63*). Raw read counts per gene were extracted using HTSeq-count v0.11.2(*64*). Gene expression profiles were constructed for differential expression, cluster, and principle component analyses with the DESeq2 package (https://bioconductor.org/packages/release/bioc/html/DESeq2.html)(*65*). For differential expression analysis, pairwise comparisons between two or more groups using parametric tests where read counts follow a negative binomial distribution with a gene-specific dispersion parameter. Corrected p-values were calculated using the Benjamini-Hochberg method to adjust for multiple testing. Gene enrichment analysis was done using Mycobrowser, BioCyc, and TBDB databases.

**Quantitative Real-Time PCR**

Reverse transcription of total RNA into complementary DNA (cDNA) was done using 500 ng RNA, 50 U/µl M-MuLV (NEB M0253), GeneAmp™ 10x PCR gold buffer (Applied Biosystems 4306898), and 50 µM random hexamers (Applied Biosystems N8080127). Residual RNA was cleared by alkaline hydrolysis and pure cDNA was concentrated using a PCR cleanup kit (Qiagen 28104). Gene expression levels were quantified using SYBR® Green PCR Master Mix (Applied Biosciences 4309155) and gene-specific primers on the LightCycler® 480 System (Roche 05015243001). Results were normalized to 16S rRNA (for Mtb) or *groEL* (for non-Mtb) and $^{\Delta\Delta}$Ct was calculated and plotted on GraphPad Prism. PrimerQuest tool from IDT Technologies (https://www.idtdna.com/PrimerQuest/Home/Index) was used to design all primers.

**Microfluidic Control of Antibiotic Exposure**

The *M. smegmatis* mc2-155 reporter strain expressing msfGFP-tagged RpoB from its genomic locus and mCherry2B-tagged FtsZ from the Tweety phage integration site was kindly provided by Dr. Cara Boutte at the University of Texas at Arlington and was maintained in 7H9-ADC (7H9 liquid media with 5 gm/liter albumin, 2 gm/liter dextrose, 0.85 gm/liter NaCl, 0.003 gm/liter catalase, 0.2% (v/v) glycerol and 0.05% (v/v) Tween 80) supplemented with 0.025 gm/liter Zeocin (Invitrogen) and 0.05 gm/liter Hygromycin B (VWR) unless specified otherwise. Microfluidic control of antibiotic pulsing (Fig. 3D and Fig. S13) was performed using the CellASIC ONIX2 system (EMD Millipore) with the B04A microfluidic plate, which was designed to accommodate bacterial samples. The reporter strain was first cultured with Zeocin and Hygromycin to reach OD600≈1.0, then diluted by ~200 fold in plain 7H9-ADC media until it reached OD600≈0.5. This exponentially growing, antibiotic-naïve *M. smegmatis* culture was further diluted to a final OD600≈0.1 and seeded into the sample loading wells of the microfluidic plate. Cells were injected into imaging flow wells by transient flow pulses and maintained at 37°C with a constant flow of plain or drug-containing 7H9-ADC liquid media.

**Time-lapse Microscopy and Image Analysis**

Time-lapse microscopy imaging was performed using a widefield Nikon Eclipse Ti-E inverted microscope equipped with Perfect Focus System for maintenance of focus over time and NIS Elements software (v. 4.5) for multi-dimensional image acquisition. The temperature of the imaging stage was maintained at 37°C using an Okolab Cage incubator. Phase-contrast and epifluorescence signals were collected every 15 minutes from a Nikon CFI Plan Apo DM Lambda ×100 1.45 NA oil objective and using with Andor Zyla 4.2 sCMOS camera. FtsZ-mCherry2B and RpoB-msfGFP were excited with a Lumencor Spectra X light engine at 100% LED power with Chroma FITC (470/24) and mCherry (575/25) filter sets, respectively, and collected with a Spectra Sedat Quad filter cube ET435/26M-25 ET515/30M-25 ET595/40M-25 ET705/72M-25 (msfGFP) and a Spectra CFP/YFP/mCherry filter cube ET475/20M-25 ET540/21M-25 ET632/60M-25 (for mCherry2B). The LED fluorescence illumination power was set to 100% and the exposure times for imaging FtsZ-mCherry2B and RpoB-msfGFP were set to 300ms and 100ms, respectively. Time-lapse microscopy data were analyzed using custom Python scripts. Briefly, planar drift between successive frames was detected via a phase-correlation method. This drift vector was used to generate a Fourier-shifted image in frequency space, which was then converted to a drift-corrected image using inverse Fourier transformation. The re-aligned large image sequences were then cropped into smaller patches using the ROI tool in Fiji. Semantic image segmentation of phase-contrast images was performed using a custom-trained UNet model. Midline coordinates of immobilized cells were generated using our previously published Python package, MOMIA (Mycobacteria-Optimized Microscopy Microscopy Image Analysis). For imaging fields where cells were not properly immobilized, we leveraged the “Segmented line” tool in Fiji to generate midline coordinates.

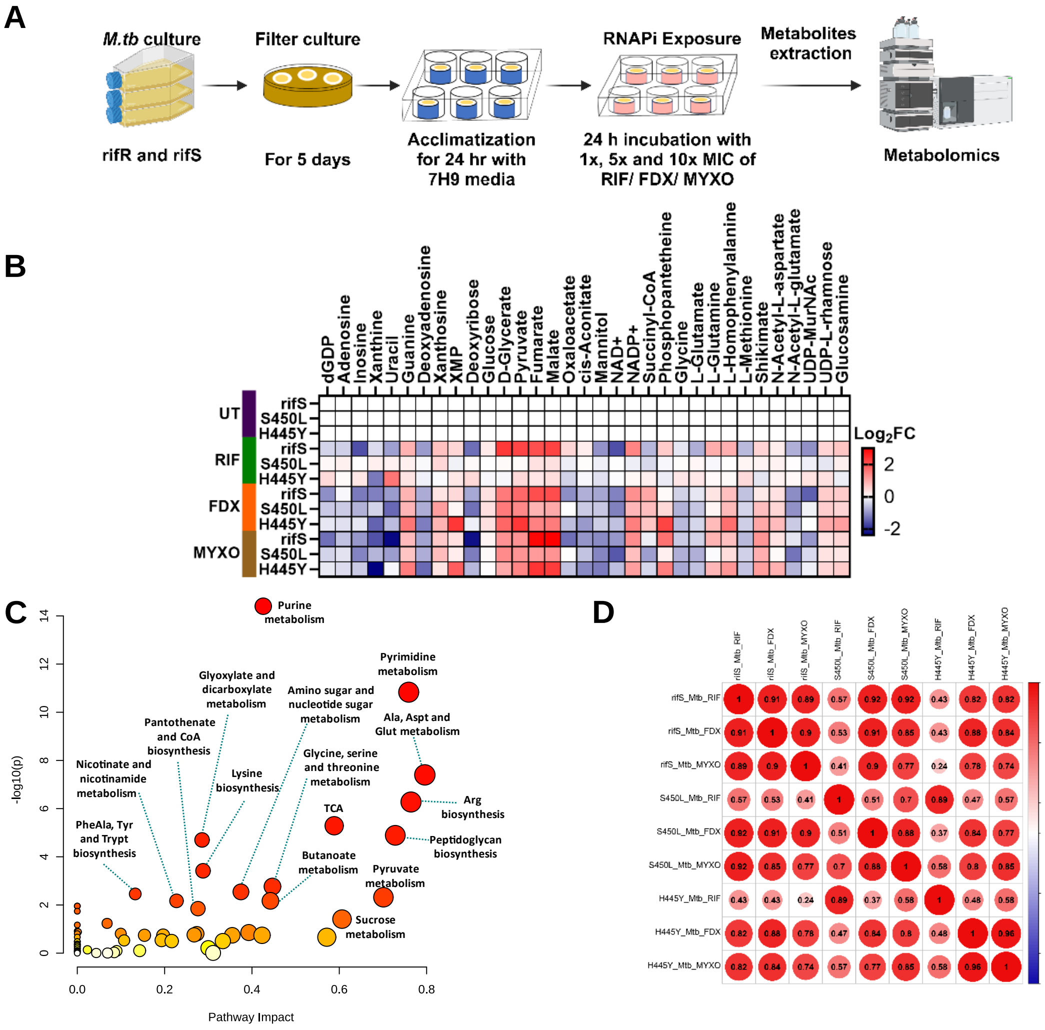


**Figure S1: Metabolic features of rifS and rifR strains of Mtb Erdman: (A)** Outline of metabolomics experiment. Mtb (rifS and rifR) strains were grown up to 0.8 $OD_{580}$ and then passed through 0.22µm filters and cultured for 5 days at 37°C on 7H10 agar plates. These bacteria-laden filters were exposed to 1x, 5x, and 10x MIC concentrations of RNAP inhibitors (RIF, FDX, and MYXO) for 24 hrs and collected for metabolite extraction followed by metabolic profiling on LC-MS. **(B)** Heatmap showing extended metabolic profile upon RNAP inhibition by RIF, FDX and MYXO. These metabolites represent purine catabolism, central carbon metabolism, and other peptidoglycan biosynthesis-related precursors. **(C)** Bubble plot representing the pathway enrichment analysis using "Metaboanalyst" server(*66*). 'X' axis is the pathway impact values from the pathway topology analysis and 'Y' axis scales the p-values from the pathway enrichment analysis. (Pathway impact = Sum of the importance measures of the matched metabolites / Sum of the importance measures of all metabolites in each pathway). Node color (varying from yellow to red) means different levels of significance of metabolites. **(D)** Correlation coefficient plot displaying Pearson correlation values (with 0.05 p-significant level) using SRplot server(*67*). Correlation value 1 shows maximum overlaps among groups. All treatments showed significant correlations except rifR strains (S450L and H445Y) treated with RIF. All results are representation of biological triplicate and two independent experiments.

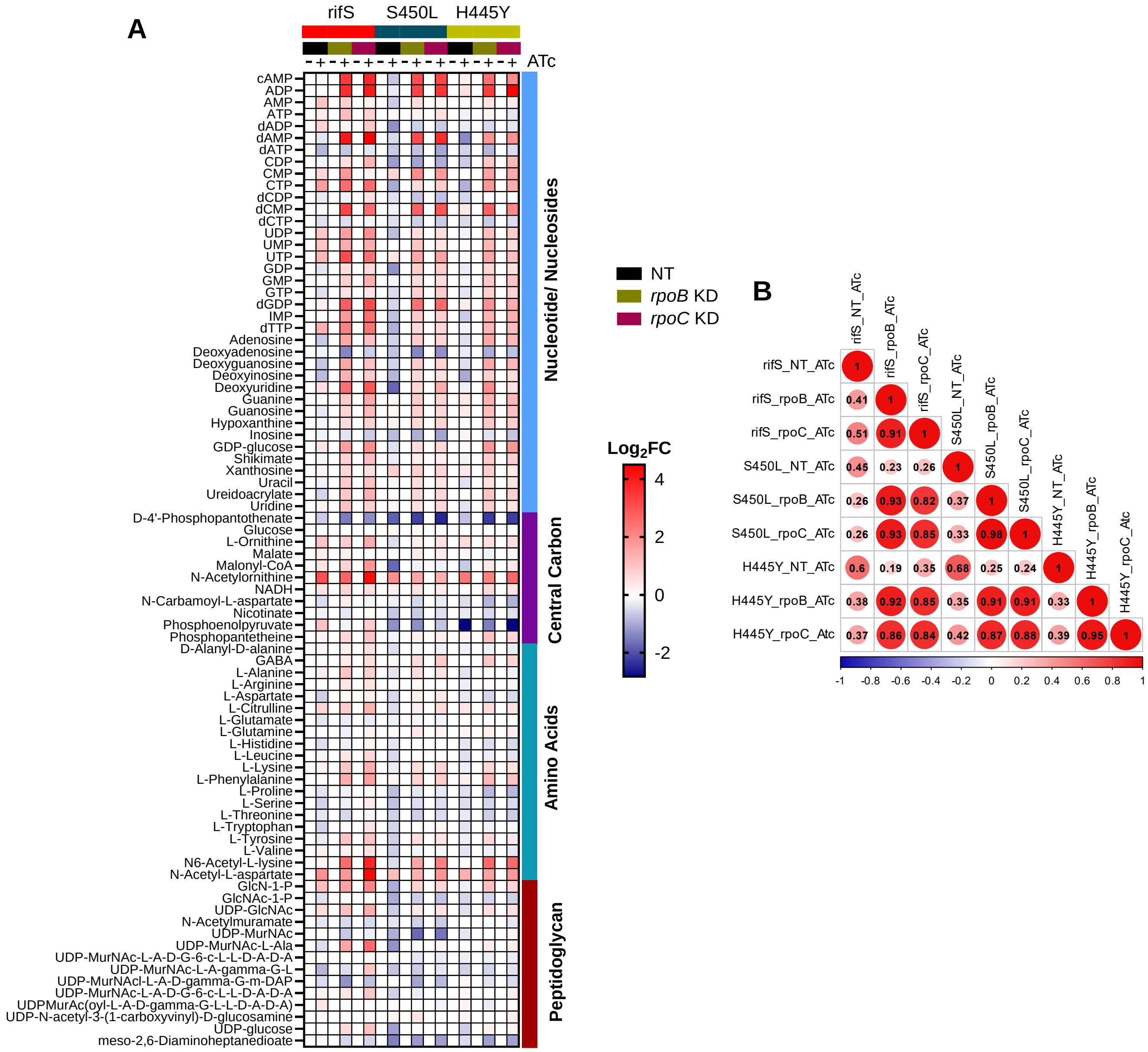


**Figure S2: Targeted metabolic analysis of CRISPRi-mediated *rpoB* and *rpoC* gene knockdowns in Mtb's RNAP complex. (A)** Heatmap illustrating Log2 fold changes in metabolite levels across various pathways following CRISPRi-mediated knockdown (KD) of RNAP subunits. **(B)** Pearson correlation coefficient plot showing the relationship among *rpoB* and *rpoC* knockdown strains on rifS and rifR strain background. *rpoB* and *rpoC* depletion leads to comparable metabolic profiles, indicating a strong correlation across different strains. All results are representation of biological triplicate and two independent experiments.

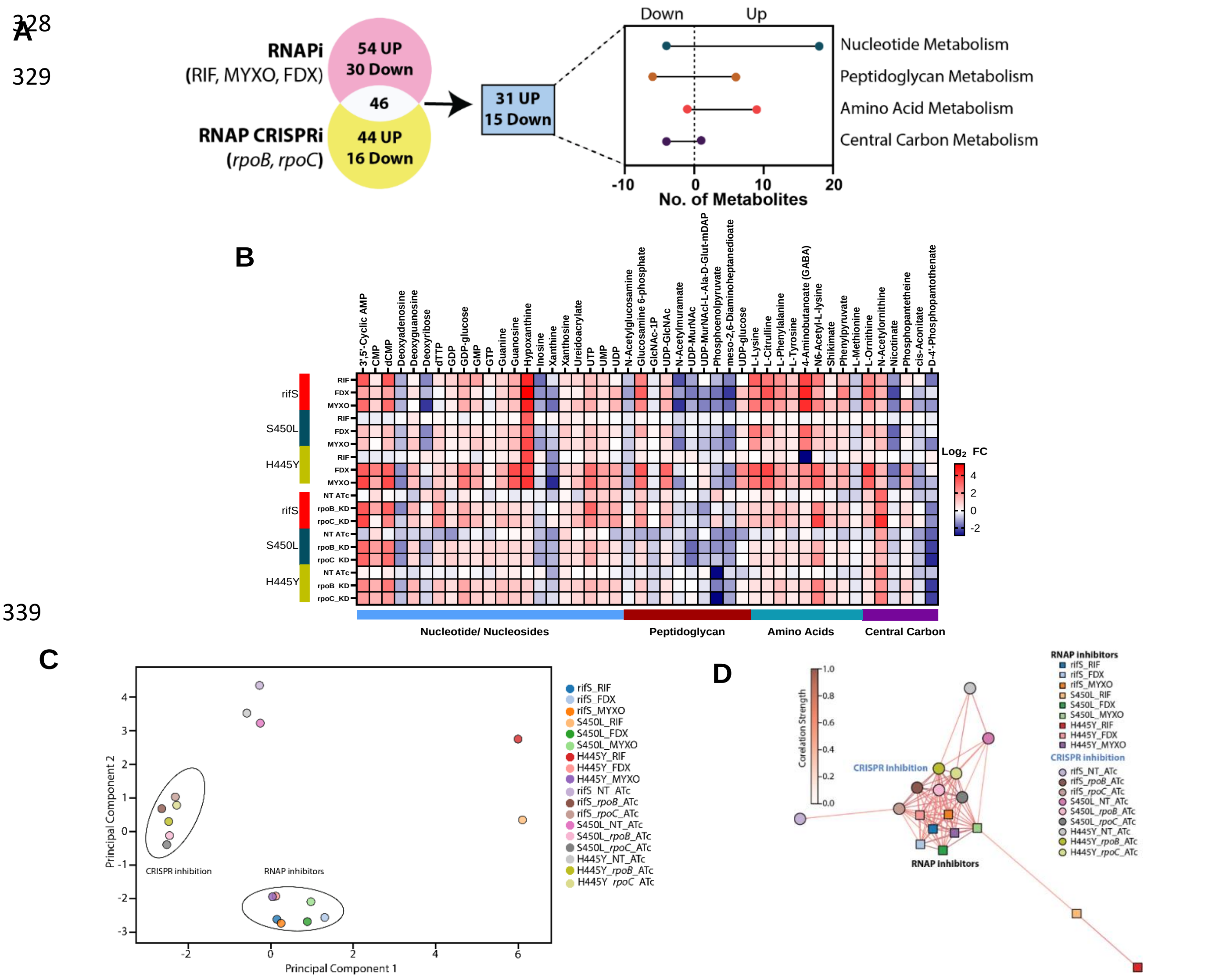


**Figure S3: Comparative analysis of metabolic profile of Mtb treated with RNAP inhibitors (RNAPi) and CRISPR mediated inhibition (CRISPRi) of *rpoB* and *rpoC* gene expression: (A)** Venn diagram showing 46 overlapping metabolites between RNAPi and CRISPRi groups with a significant overrepresentation from peptidoglycan biosynthesis. **(B)** Heatmap presenting common 46 metabolites showing similar changes ($Log_2FC$) between RNAPi and CRISPRi group. **(C)** PCA plot showing clustering of RNAPi and CRISPRi groups. **(D)** Network plot representing the significant correlation between two groups using 46 overlapping metabolites (i) Mtb treated with RNAP inhibitors (RNAPi) such as RIF, FDX, and MYXO (ii) Mtb strains with CRISPRi inhibition (CRISPRi) of *rpoB* and *rpoC*. Each node symbolizes a different treatment (squares) or gene inhibition (circles) in rifS and rifR strains. Person correlation and threshold (k) values were used for the clustering. The "length" or distance between nodes is determined by the "spring_layout" algorithm, which positions nodes closer if they have stronger connections (thicker and darker lines) and further apart if their connection is weaker or nonexistent (non-targeting, NT or rifR strains treated with RIF). Tightly grouped nodes clustered in the center denote RNAP-specific profiles that are highly correlated with each other. The colors of the lines connecting the nodes represent the correlation scores, as shown by the gradient color scale.

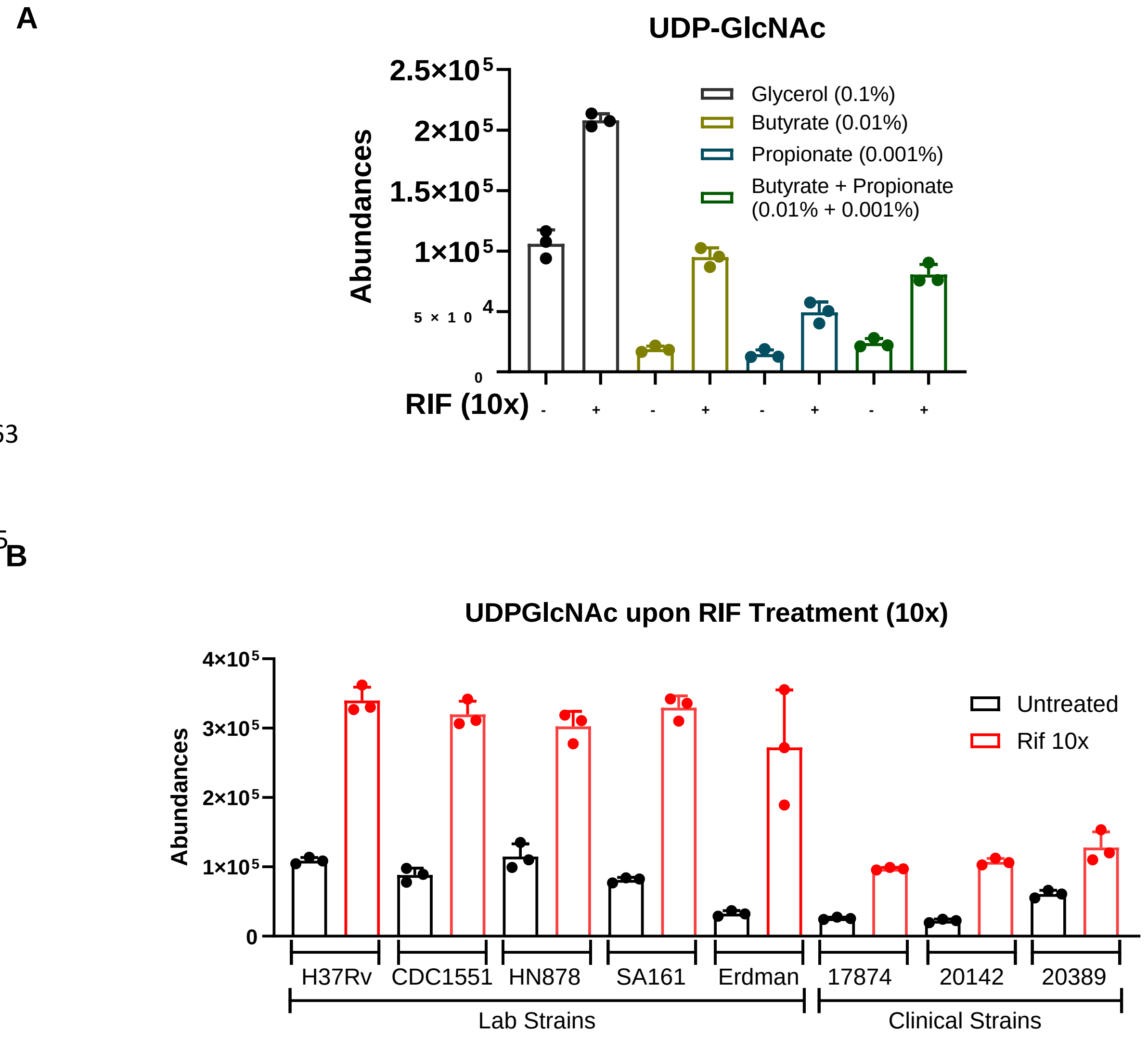


**Figure S4: UDP-GlcNAc levels (A)** in Mtb-H37Rv grown on different carbon sources (solely) and treated with 10x MIC dose of RIF for 24 hr at 37°C. 'Y' axis represents the total abundances of the respective ions. **(B)** in different laboratory-adapted and clinical isolates of Mtb treated with 10x MIC dose of RIF (0.5 µg/ml) for 24 hr at 37°C.

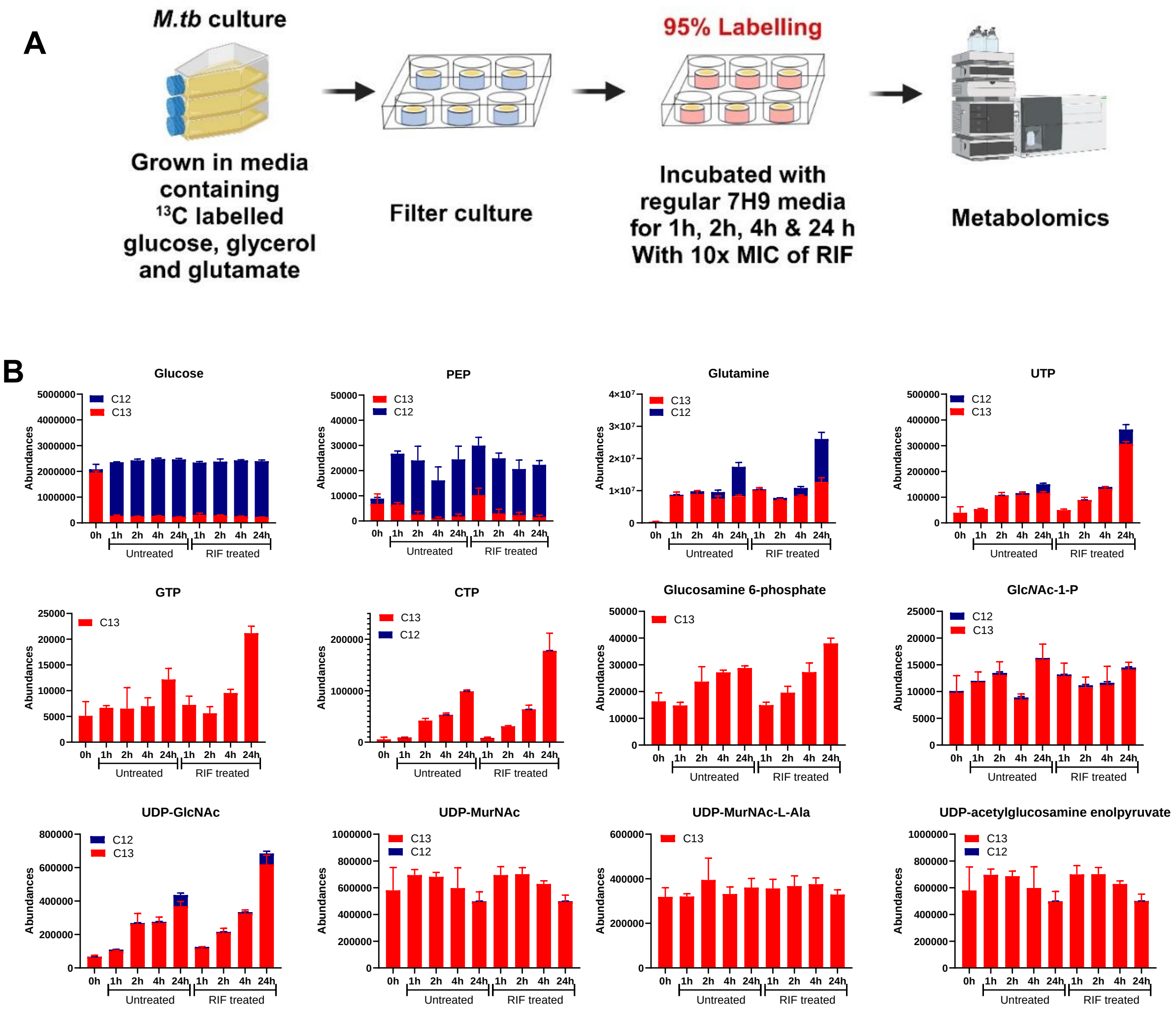


**Figure S5: Metabolic labeling of Mtb-H37Rv using $^{13}$C labeled carbon sources. (A)** Mtb was grown on modified 7H9 media with $^{13}$C labeled glucose, glycerol, and glutamate. After achieving 95% labeling, Mtb-laden filters were exposed to unlabeled defined 7H9 media with 10x MIC dose of RIF (0.5 µg/ml). Samples were collected at 1h, 2h, 4h, and 24h for metabolomic profiling. **(B)** Stack bar graphs represent the abundances of different metabolites in RIF-treated and untreated samples. Red bars denote the labeling while blue bars show the unlabeled metabolite pools. Untargeted analysis reveals accumulation of UTP, GTP, CTP, and UDP-GlcNAc in RIF-treated Mtb, as indicated by minimal replacement of labeled metabolites with unlabeled ones. All results are a representation of biological triplicate and two independent experiments.

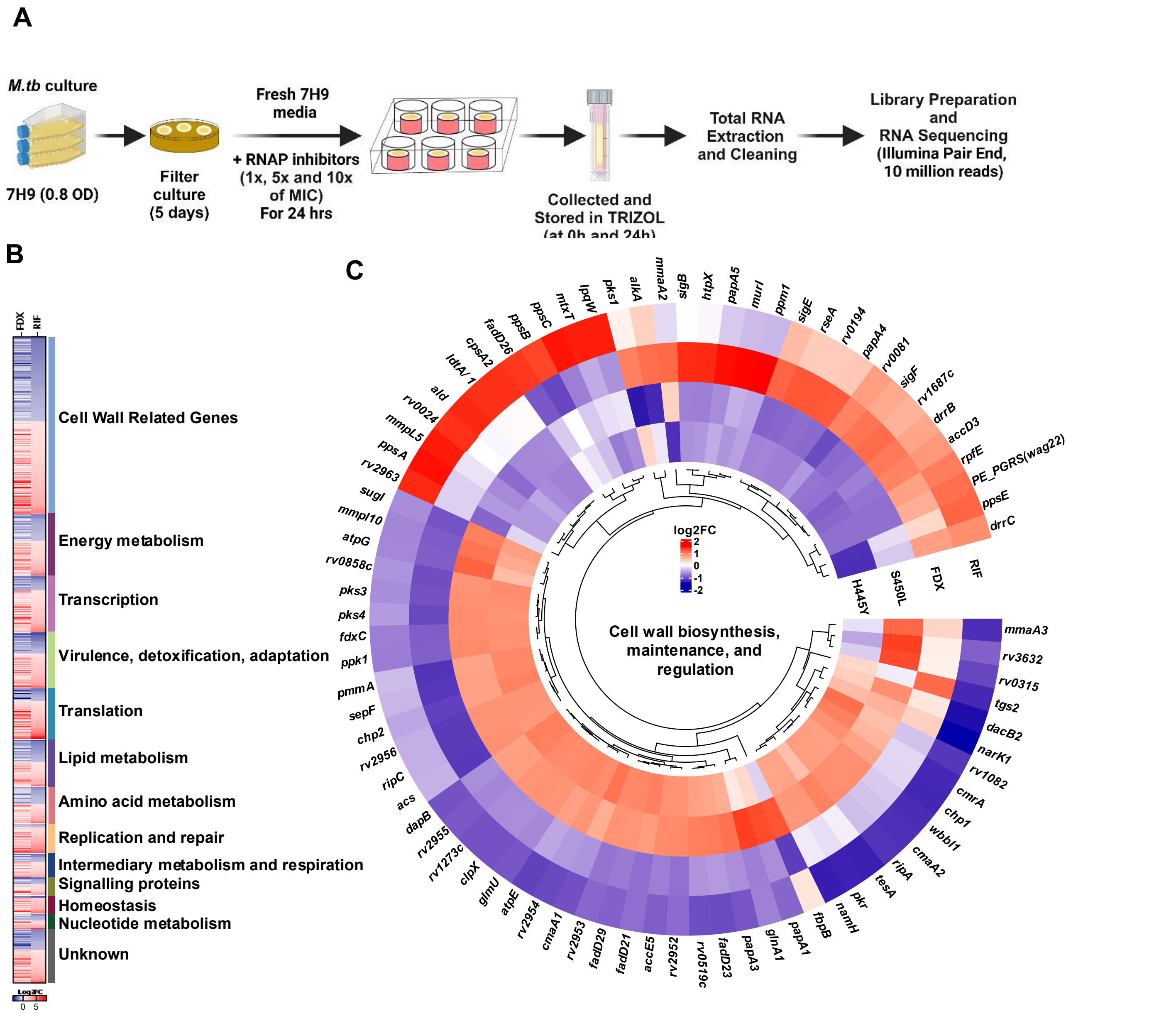


**Figure S6: Transcriptomic profiling of Mtb_H37Rv after RIF and FDX treatments. (A)** Mtb-laden filters were prepared from 0.8 $OD_{580}$ culture followed by 24h preadaptation on 7H9 and then exposure to 1x, 5x, and 10x MIC doses of rifampicin (RIF) and fidaxomicin (FDX) for the next 24h. Samples were collected in GTC buffer and stored in TRIZOL to extract total RNA. mRNA sequencing was done using Illumina's Pair End methods with 10 million reads. RifS strains were exposed to RIF and FDX while rifR strains were treated with RIF alone. **(B)** Heatmap showing the expression pattern of genes from different cellular functions. Red-blue color scale is used to represent statistically significant $\log_2$ fold changes. Gene enrichment and classification was done using Mycobrowser, BioCyc, and TBDB databases. **(C)** Circular cluster heatmaps showing $\log_2$ fold changes of genes annotated to function in cell wall maintenance, biosynthesis, regulation, and cell division in red-blue color scale. Clusters are calculated using 'complete' clustering method while Euclidean distance method was used to plot the distance tree. All results are average of biological triplicates.

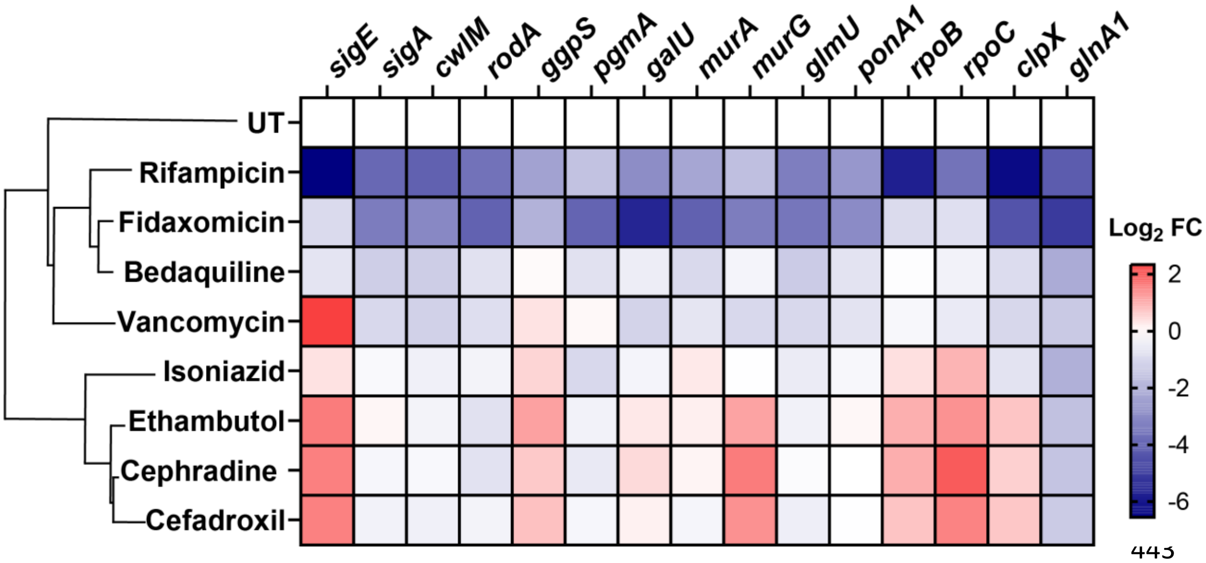


**Figure S7: qPCR analysis of genes related to peptidoglycan biosynthesis.** Mtb-H37Rv-laden filters were treated with 10x MIC doses of RNAP-targeting antibiotics (RIF and FDX), cell wall targeting antibiotics (ethambutol, isoniazid, cephradine, cefadroxil, and vancomycin) and bedaquiline (anti-respiratory) for 24h at 37°C. Total RNA was extracted followed by cDNA preparation. Gene levels (log$_2$fold change) were estimated by qPCR using gene-specific primers.

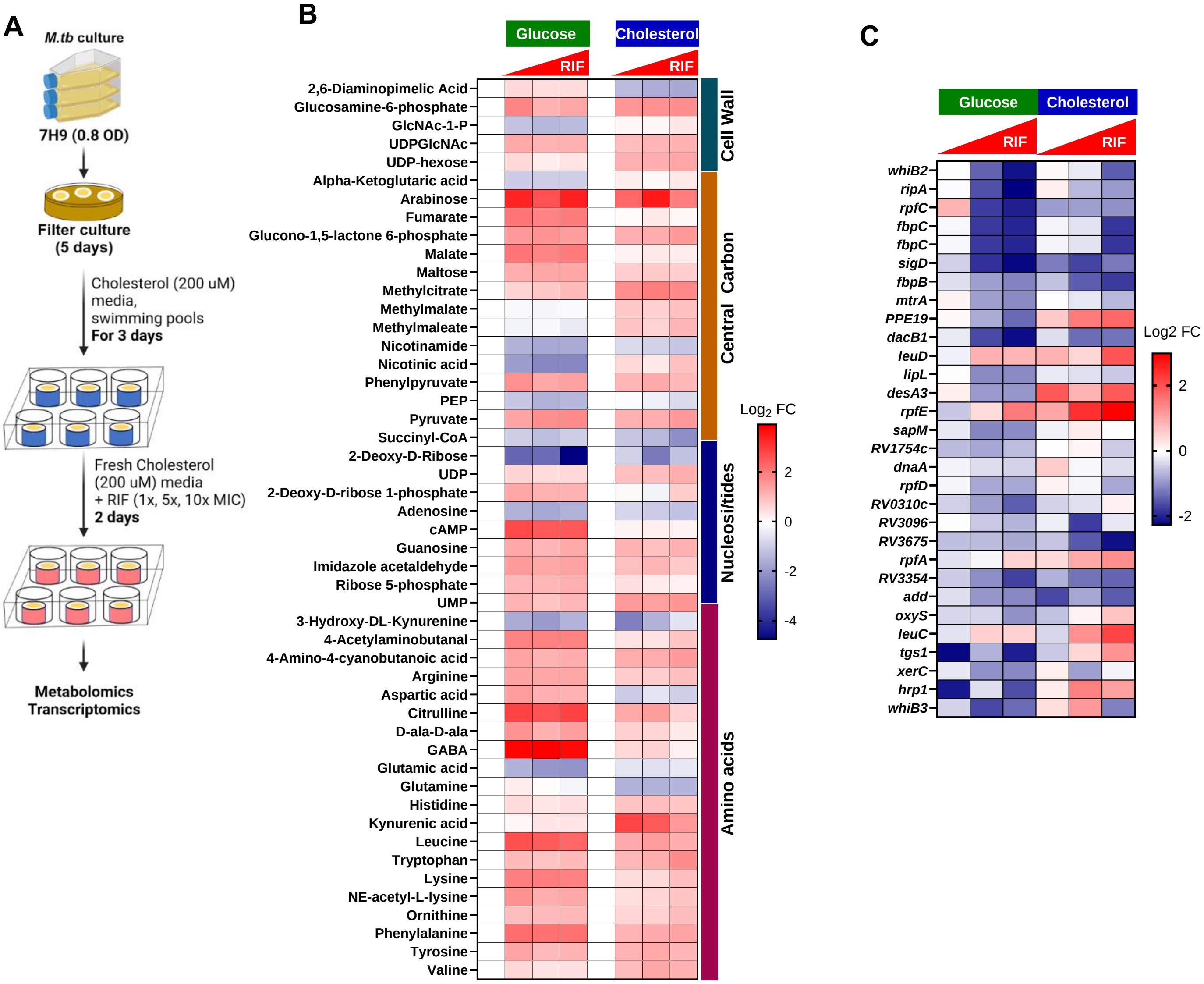


**Figure S8: Association of RNAP perturbation with PG biosynthesis and *mtrAB* regulon expression are carbon source independent. (A)** 1 ml cultures of Mtb (0.8 $OD_{580}$) were grown on filters for 5 days followed by acclimatizing on cholesterol or glucose-containing modified 7H9 media for 3 days (for cholesterol) or 24 hr (glucose). Filters were exposed to different doses of RIF (1x, 5x, and 10x) for 48h (cholesterol) and 24 hr (glucose) followed by transcriptomic and metabolomic profiling. **(B)** Metabolomic profile upon RIF treatment shows significant overlaps between cholesterol and glucose grown Mtb. **(C)** Heatmap showing carbon source independent downregulation of *mtrAB* regulon genes.

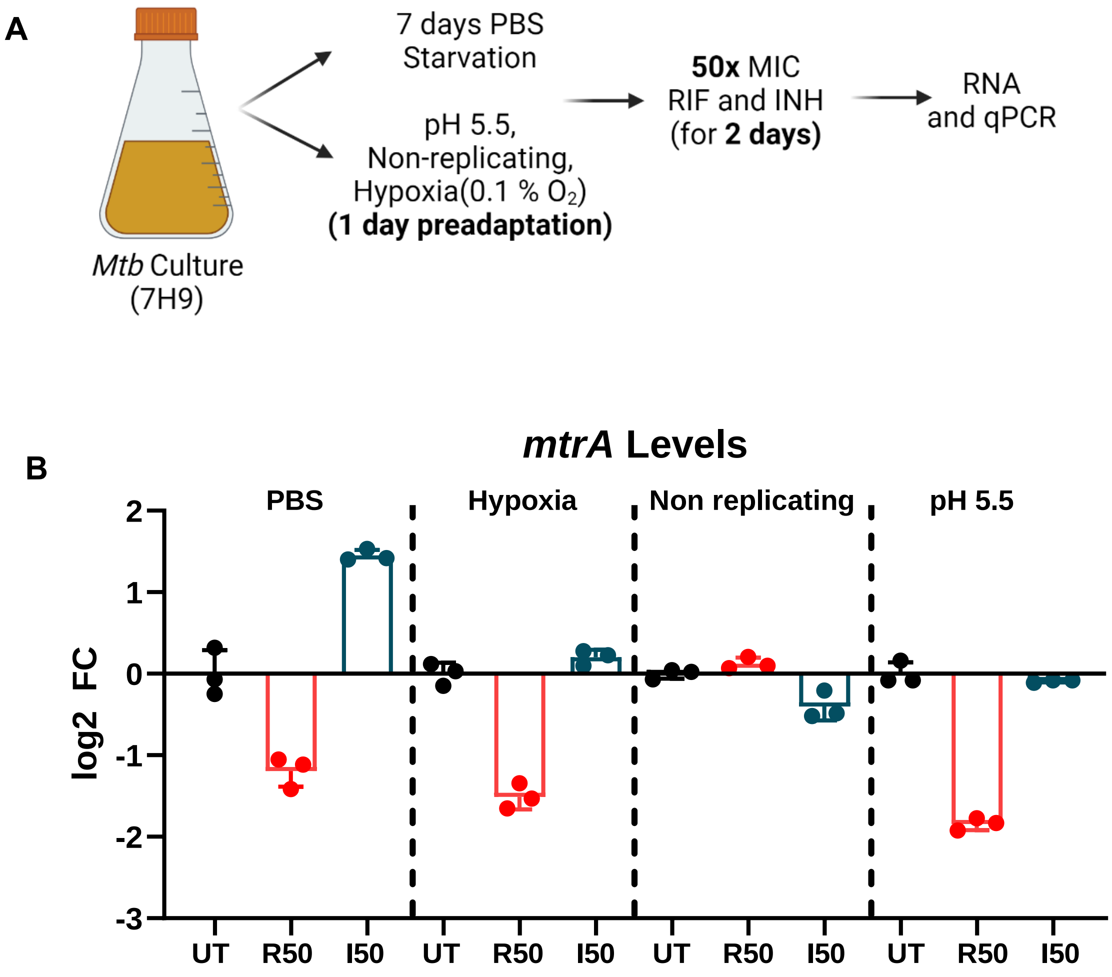


**Figure S9: *mtrA* levels upon stress exposure and treatment with RIF and INH. (A)** 0.8 $OD_{580}$ Mtb-Erdman cultures were preadapted to stress conditions (PBS starvation for 7 days, and 1 day for pH 5.5, non-replicating media, and hypoxia) before being treated with vehicle or 50x MIC RIF and INH for 2 days. Total RNA was then extracted, and *mtrA* expression was quantified by qPCR. **(B)** Bar graph representing the log2 fold changes of *mtrA* expression with respect to untreated samples. UT= untreated; R50= rifampin; I50= isoniazid.

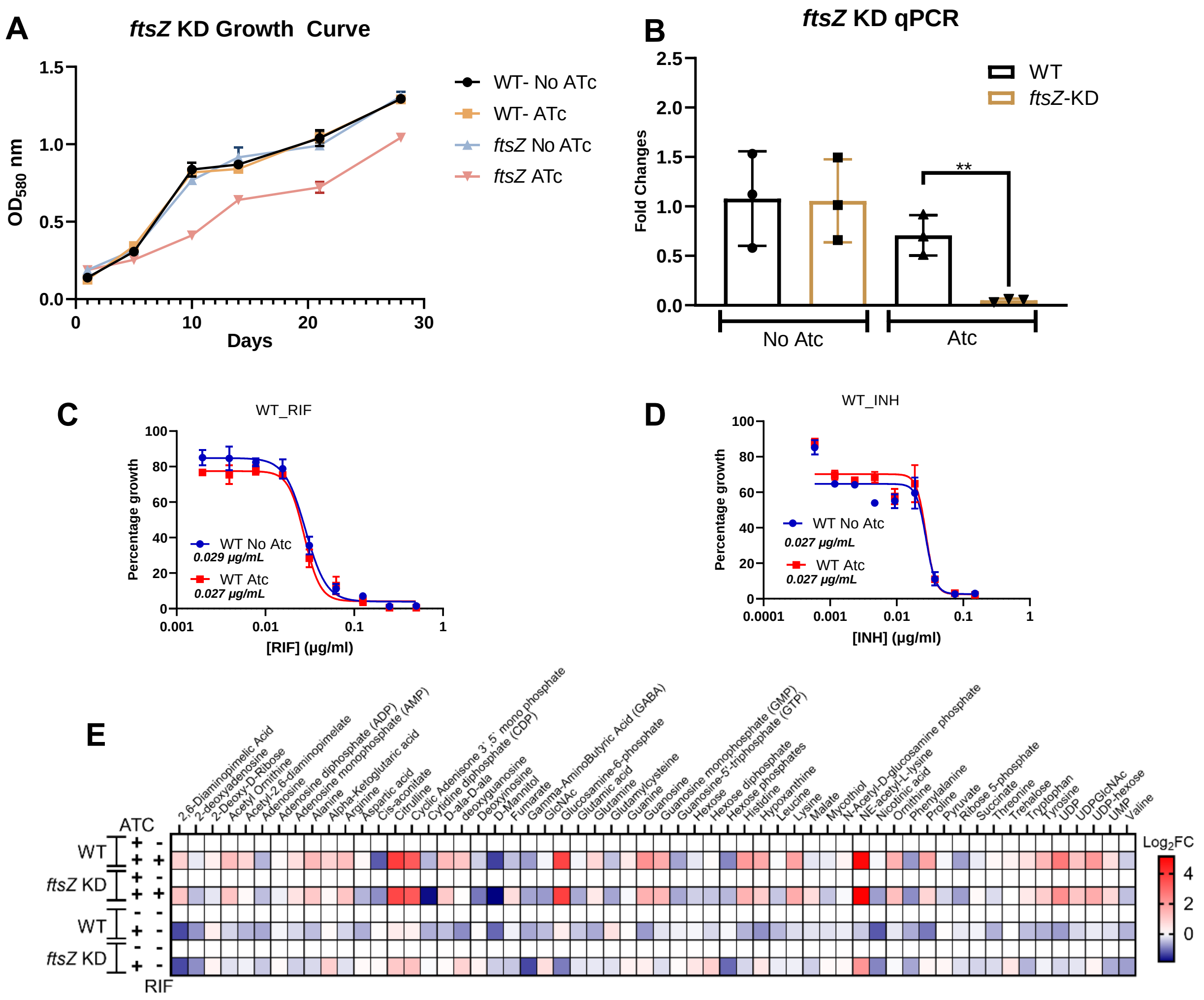


**Figure S10: CRISPRi depletion of *ftsZ* in Mtb-H37Rv. (A)** Growth curve shows controlled reduction in growth upon *ftsZ* silencing. **(B)** Planktonic cultures of wild type (WT) and CRISPRi gene depletion mutant of *ftsZ* were treated with 100 nM of ATc for 8 days and collected to extract total RNA and qPCR analysis. Results suggest a significant reduction in *ftsZ* expression in *ftsZ* KD. **(C-D)** Dose-response curves for WT strains treated with RIF and INH and with and without ATc, were generated to assess the impact of ATc on antibiotic susceptibility. **(E)** Heatmap presenting the metabolic profile of WT and *ftsZ* KD strains both in the presence and absence of RIF and ATc. RIF treatment was done using a 10x MIC dose for 24 hr. All results are average of biological triplicates.

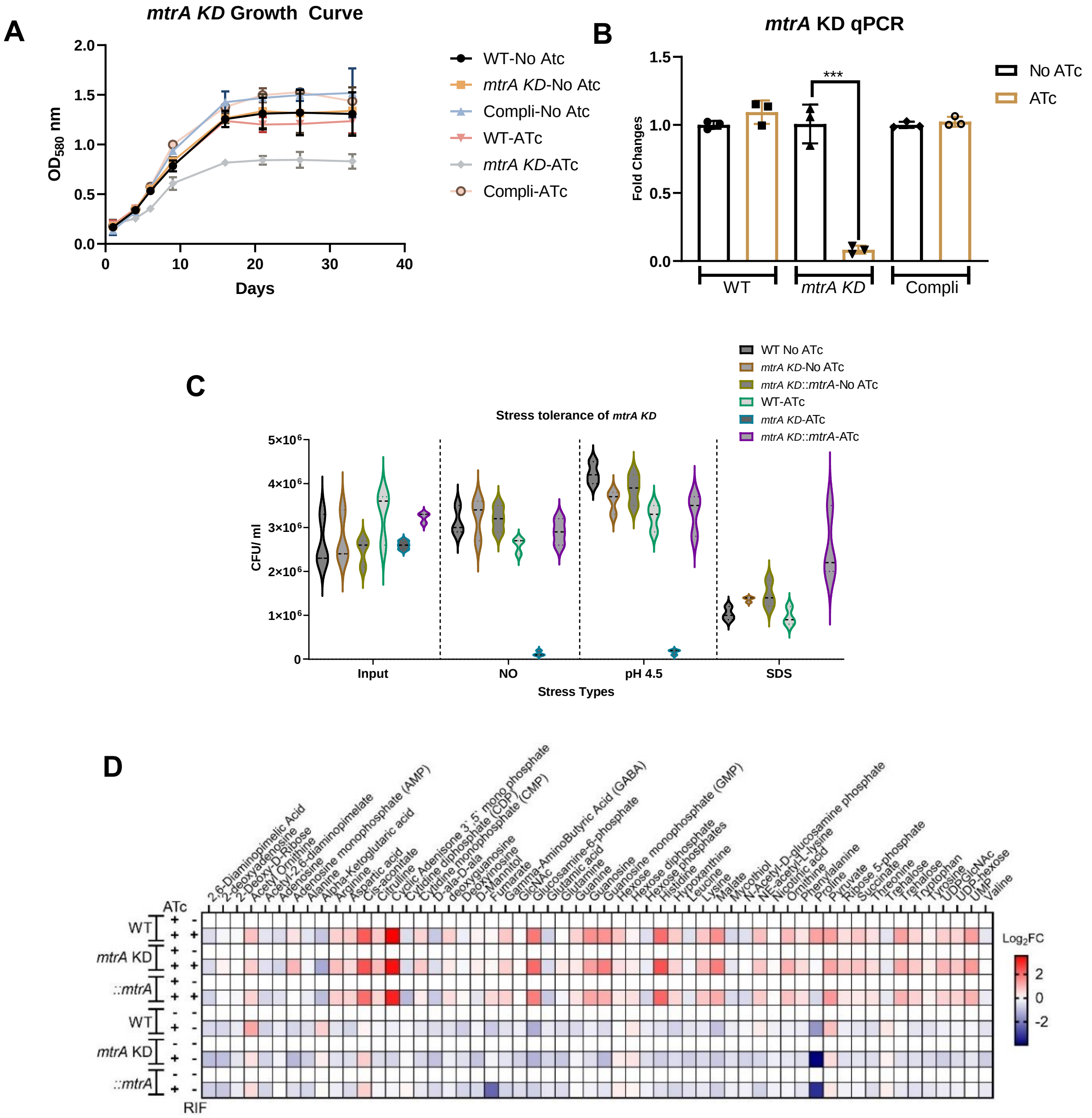


**Figure S11: Impact of *mtrA* KD on stress tolerance and metabolome of Mtb H37Rv. (A)** Growth curve demonstrating the growth pattern of WT, *mtrA* KD, and complementary (compli) strain of *mtrA* in the presence and absence of ATc. **(B)** qPCR analysis shows the fold change in *mtrA* expression after 5 days of depletion. **(C)** Violine plot showing the colony forming units (CFUs) based viability of Mtb upon *mtrA* depletion and stress exposure. All strains were exposed to pH 4.5 for 3 days, 0.1% SDS for 5 hr, and 5 mM NaNO2$^{+}$ pH=5.5 for 3 days in 7H9 media with 0.05% tyloxapol. Both input and final-day samples were processed for the CFU plating. **(D)** Heatmap showing the metabolic profile of WT, *mtrA* KD, and *mtrA* compli strains both in the presence and absence of RIF and ATc. RIF treatment used a 10x MIC dose for 24 hr.

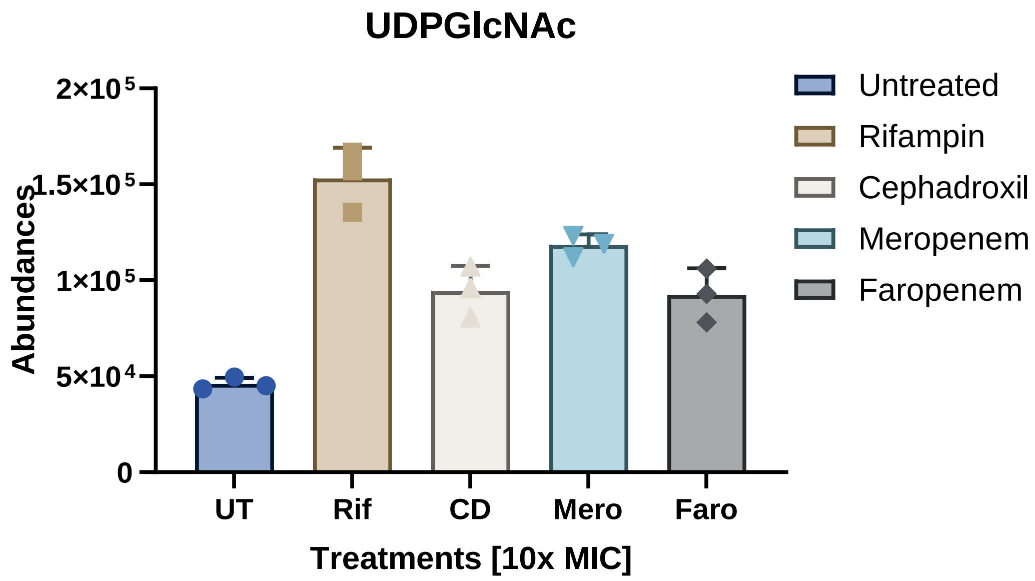


**Figure S12: UDP-GlcNAc levels upon β-lactam treatments.** Filters laden with H37Rv were treated with 10x MIC doses of various β-lactam antibiotics for 24 hours. Following treatment, metabolomic profiling was performed. Targeted analysis revealed an increase in UDP-GlcNAc levels, similar to the effect observed with RIF treatment.

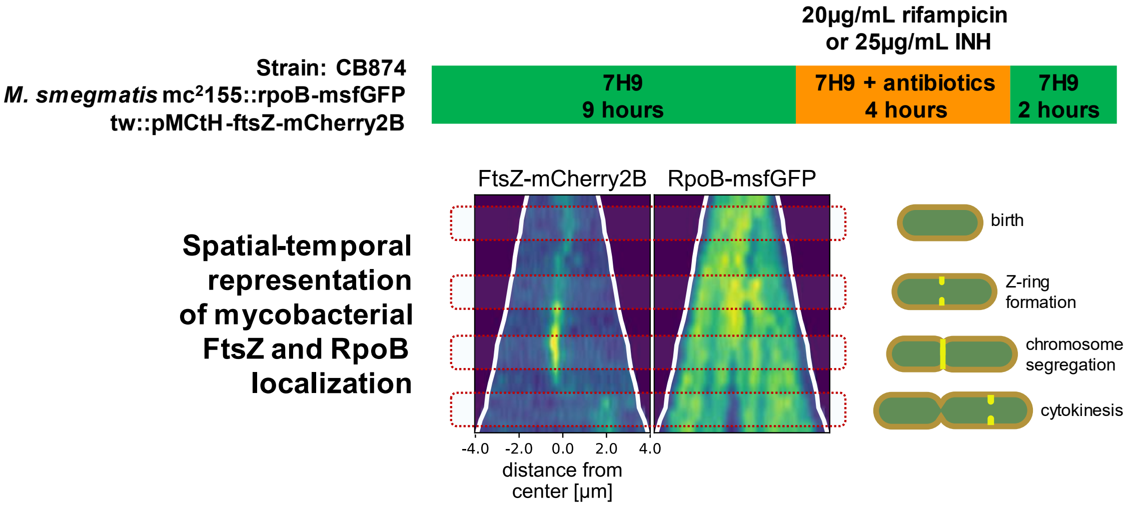


**Figure S13: Schematic representing the subcellular distribution dynamics of FtsZ and RpoB upon RIF and INH Treatment.** Recombinant *M. smegmatis* mc$^2$155 strains expressing msfGFP labeled RpoB and mCherry2B labeled FtsZ proteins were grown in a microfluidic chamber for 9h in 7H9 media followed by 4h of antibiotic treatment and 2h of recovery. Time-lapse microscopy was used to measure the distribution and expression dynamics of RpoB and FtsZ under antibiotic treatments. Fluorescence kymographs were then analyzed using MOMIA (mycobacteria-optimized microscopy microscopy image analysis). 'X' axis shows the distance from the center and the 'Y' axis represents time since the last division.

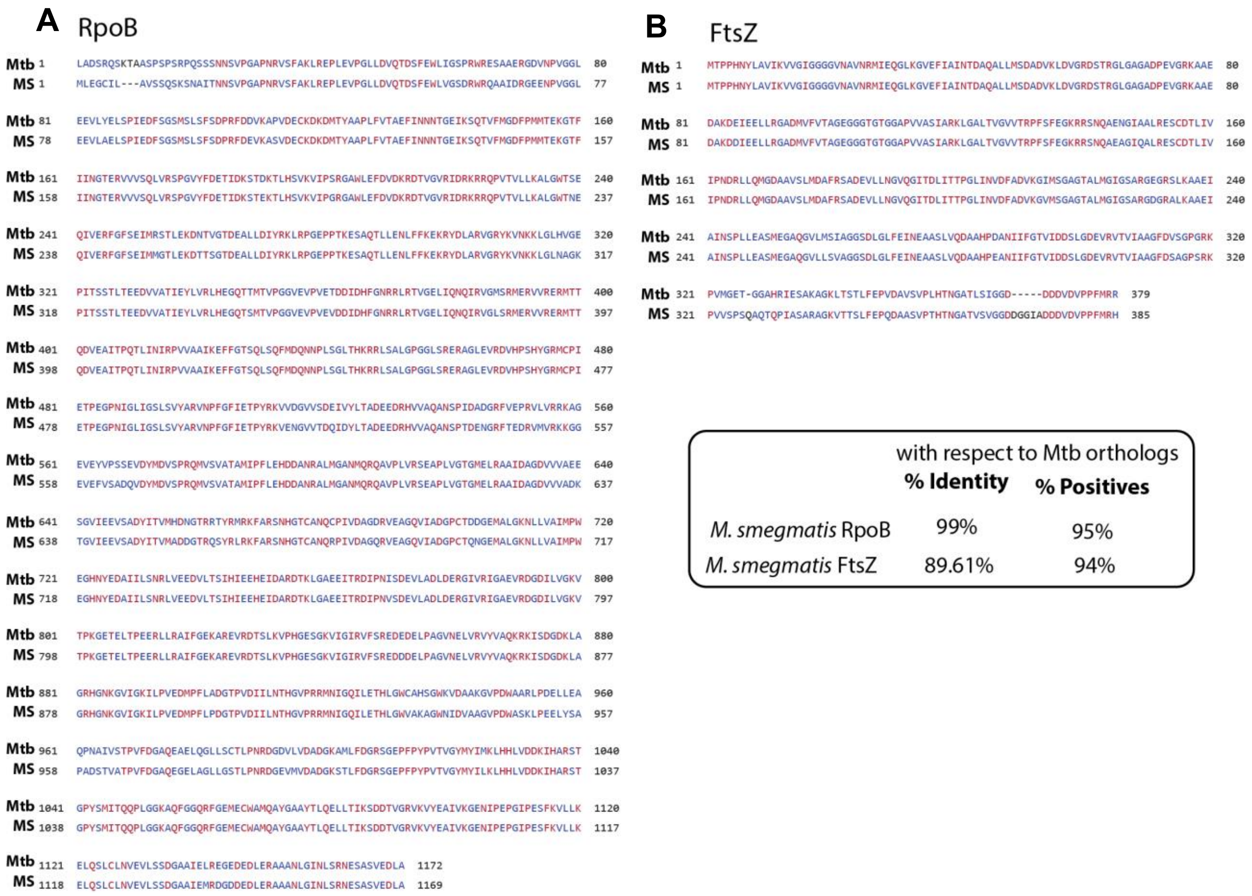


**Figure S14: Protein sequence alignments of Mtb and *M. smegmatis* (MS) RpoB and FtsZ. (A)** Mtb-RpoB protein sequence aligned with MS-RpoB showing 99% identity and 95% positives in amino acid sequences (inset). **(B)** Alignment of Mtb-FtsZ protein sequence with MS-FtsZ shows 89.61% identity and 94% positives in amino acid sequences (inset). All sequences were taken from the Mycobrowser platform and aligned using BLAST protein software.

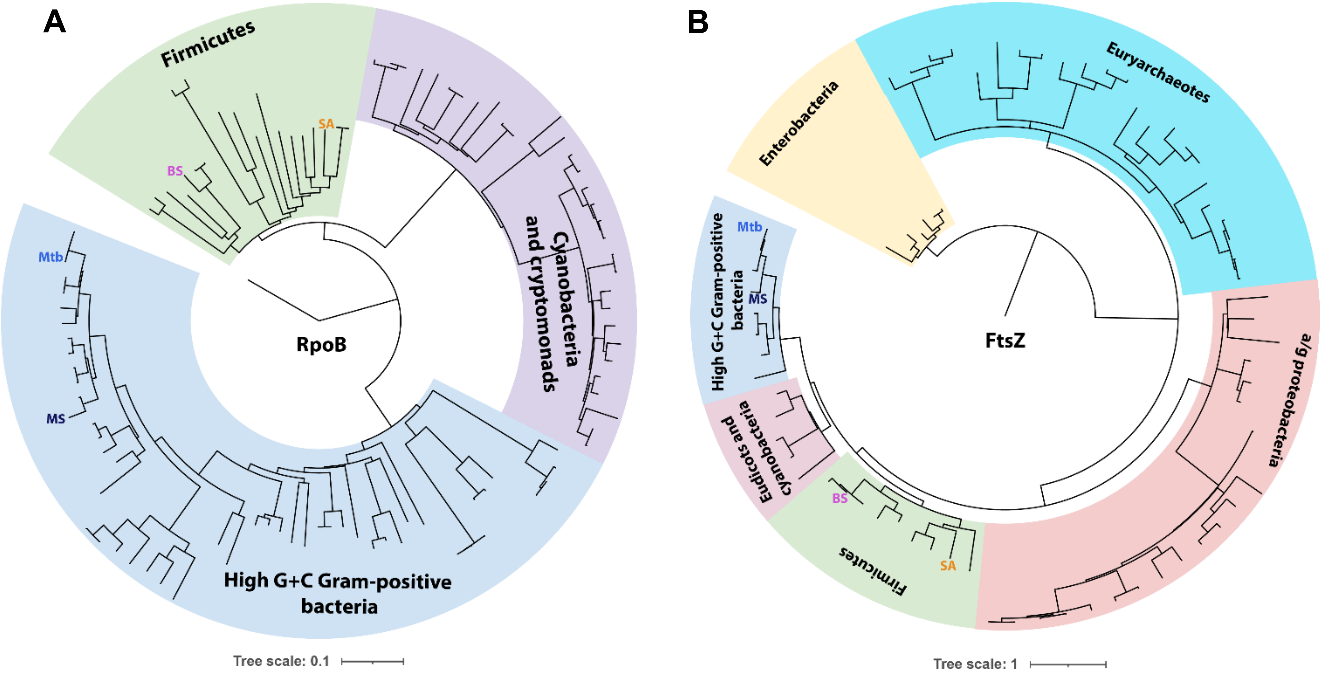


**Figure S15: Phylogenetic distribution of RpoB and FtsZ across different bacterial species. (A)** The circular dendrogram shows the distance tree results of RpoB protein across different microbial phyla. **(B)** Distance tree results of FtsZ protein from various bacterial phyla. Mtb: *M. tuberculosis*; MS: *M. smegmatis*; BS: *B. subtilis*; SA: *S. aureus*. Protein sequences were extracted from Protein-BLAST using UniProtKB/Swiss-Prot(swissprot) database. Distance tree results were exported from the BLAST tree view and visualized on https://itol.embl.de/ platform. Parameters were as follows: (i) Tree method: Fast minimum Evolution, (ii) Max Seq Difference: 0.85, (iii) Distance method: Grishin (protein).

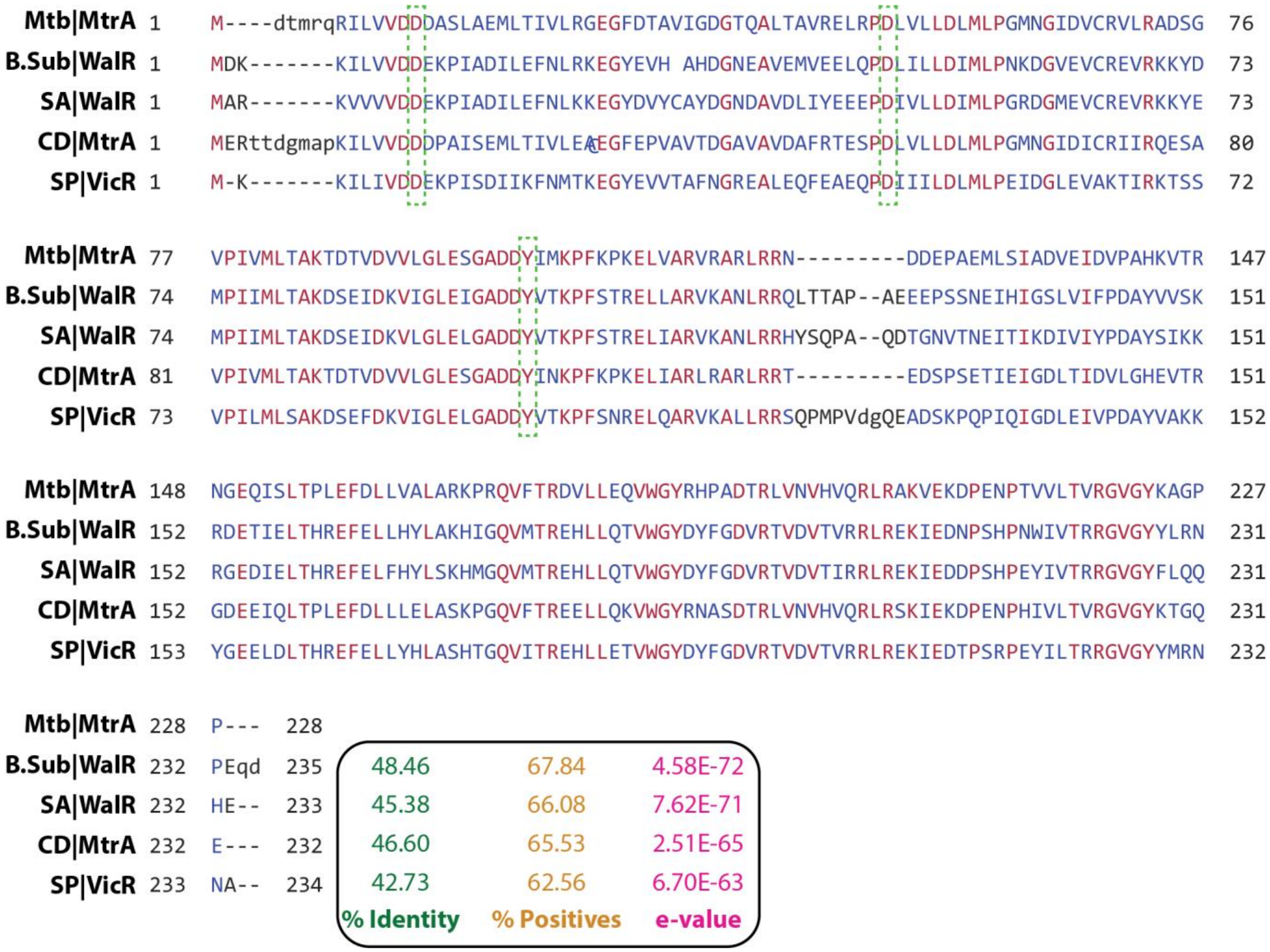


**Figure S16: Sequence alignments of MtrA orthologs:** Protein sequences of MtrA and its orthologs from *Bacillus subtilis* (B. Sub), *Staphylococcus aureus* (SA), *Corynebacterium diphtheriae* (CD) and *Streptococcus pneumoniae* (SP) were aligned using COBALT alignment tool. Inset is showing %identity, %positives, and e-values. Conserved aspartates (of N-terminal regulatory domain) and tyrosine (from C-terminal, winged helix-turn-helix DNA binding domain) are highlighted with green boxes. Aspartates are phosphorylated by sensor kinase (MtrB/ WalK/VicK/YycG) while tyrosine is known to be phosphorylated by the serine/threonine protein kinase (PknB or Prkc).

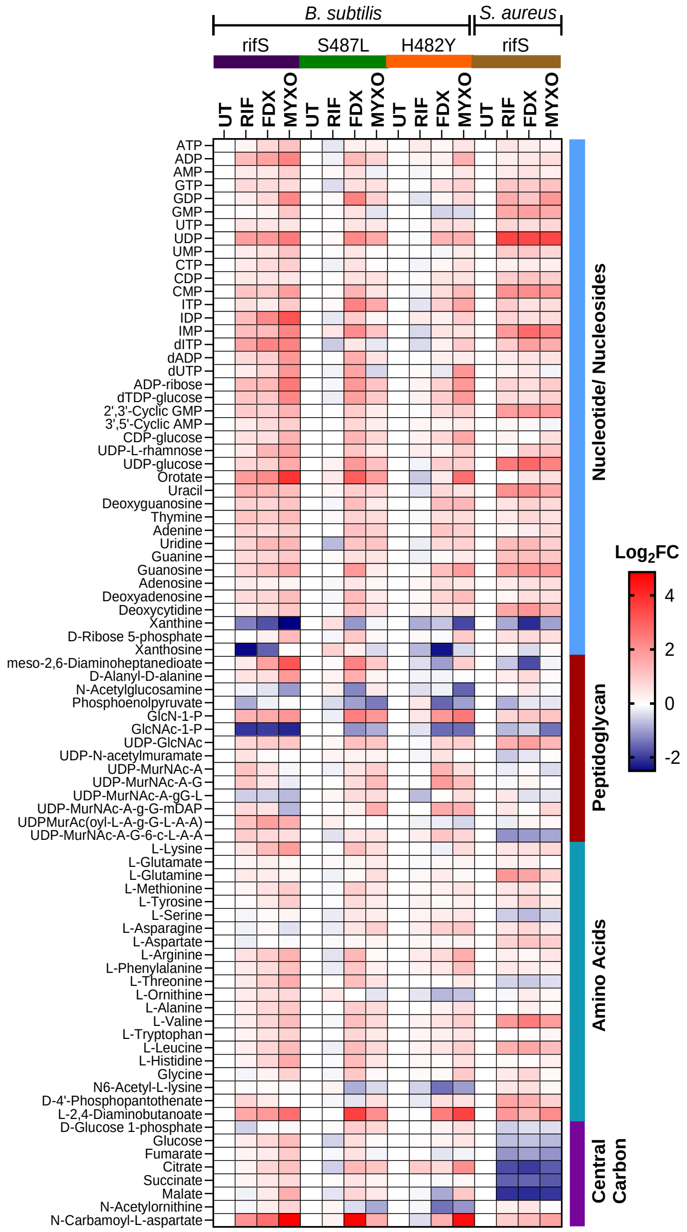


**Figure S17: Metabolomics profile of *B. subtilis* and *S. aureus* strains.** Planktonic cultures of wild type (rifS) and RIF resistant (rifR) strains (only for *B. subtilis*) were grown up to 0.45 $OD_{600}$ and treated with 10x MIC doses of different RNAP targeting drugs (RIF, FDX, MYXO), or vehicle (UT) for 20 min and collected by centrifugation (4,000 rpm, 10 min, 25°C). Pellets were washed twice with 1x PBS and resuspended in 40:40:20 (acetonitrile:methanol:water) mixture and lysed using the bead beating method. Lysates were processed and profiled on LC-MS followed by targeted analysis. Results demonstrate significant signature overlaps with Mtb treated similar RNAPi (Fig S20). All results are average of biological triplicates.

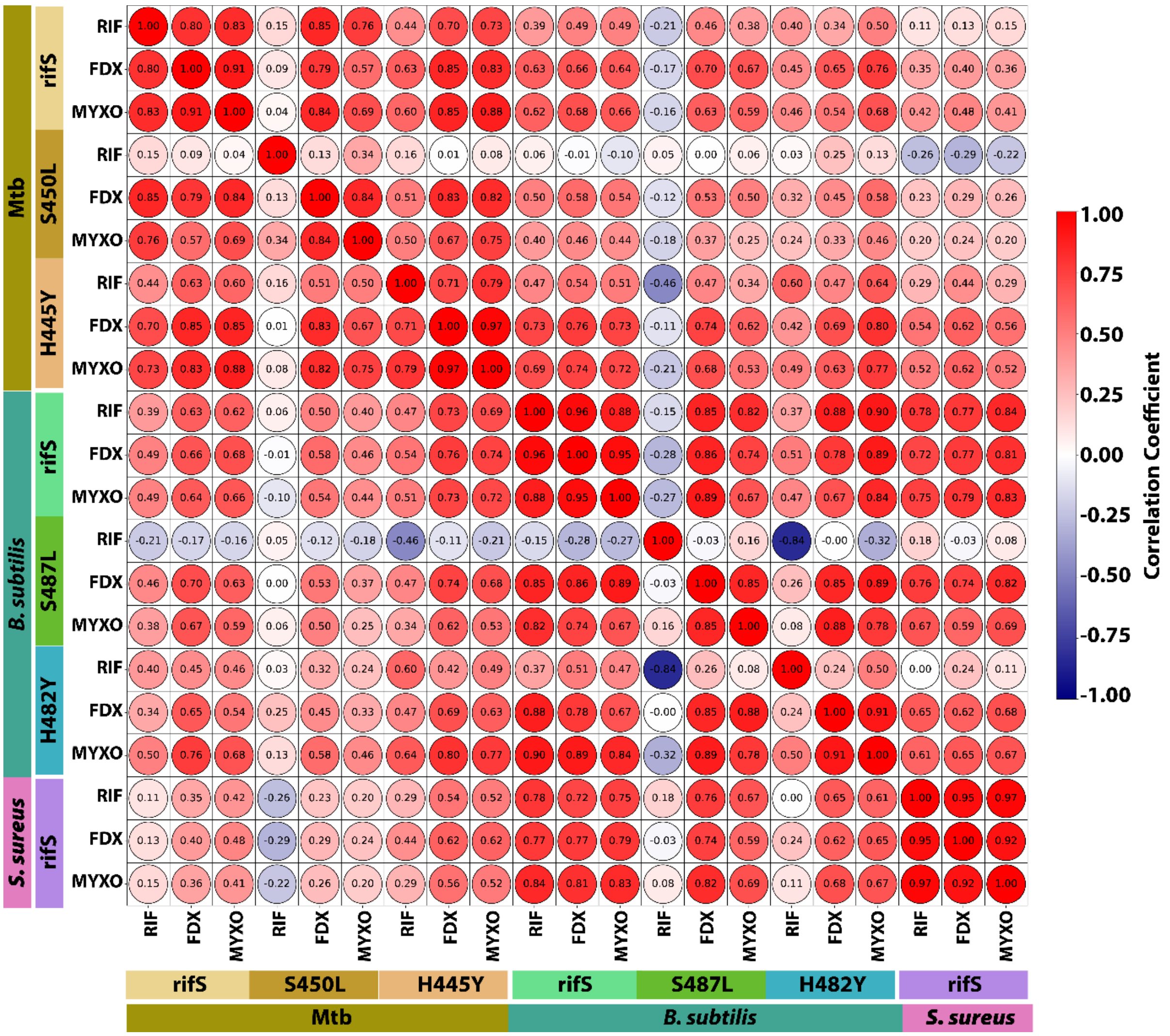


**Figure S18: Pearson correlation score plot showing conserved metabolic signature between RNAPi-treated Mtb and non-Mtb bacteria.** Key metabolic signatures were used for the analysis. Higher correlation denoted by red color or score 1. Results suggest significant overlaps between both groups.

**Table S1: Mutation in RNAP subunits (*rpoB* and *rpoC*) and its association with cell wall functions and antibiotic resistance**

| Bacteria | Physiological impacts | Reference |
|---|---|---|
| *M. tuberculosis* | Increased *dnaE2* expression (*rpoB*) | (*68*) |
| | Changed PDIM levels (*rpoB*) | (*69*) |
| | Altered colony morphology, bacillary length, cell wall thickness, permeability, and increased ofloxacin MIC (*rpoB*) | (*42*) |
| | Dysregulation in cell wall synthesis enzymes (*rpoB*) | (*44*) |
| *E. coli* | Improved osmotolerance (*rpoB*) | (*70*) |
| *B. subtilis* | Changes in competency, germination, and sporulation (*rpoB*) | (*71*) |
| | Altered levels of peptidoglycan precursors and susceptibility to β-Lactams (*rpoB*) | (*43*) |
| | Cephalosporin resistance (*rpoC*) | (*51*) |
| *S. aureus* | Better biofilm (*rpoB*) | (*72*) |
| | Heteroresistance to daptomycin and vancomycin (*rpoB*) | (*47-49*) |
| | Promotes phenotypic conversion of hVISA-to-VISA (*rpoB*) | (*73*) |
| | High-level β-lactam resistance (*rpoB, rpoC*) | (*52*) |
| *N. meningitidis* | Decreased cell membrane permeability (*rpoB*) | (*45*) |
| *Neisseria gonorrhoeae* | Cephalosporin resistance (*rpoB, rpoD*) | (*53*) |
| *Enterococcus faecalis* | Altered intrinsic cephalosporin resistance (*rpoB*) | (*54*) |

**Table S2: Associations between MtrAB and its orthologs (WalKR) with cell wall biology, antibiotic sensitivity, and resistance.**

| **Bacteria** | **Physiological impact** | **Reference** |
|---|---|---|
| *M. tuberculosis* | MtrA affects cell division, intrinsic resistance, and tolerance | (*10, 11*),(*9*) |
| | MtrB interacts with FtsZ, FtsI, and Wag31 at septa and affects antibiotic sensitivity. | (*11, 23, 24*) |
| *B. subtilis* | *walK* inhibition results in higher RIF sensitivity | (*74*) |
| *S. aureus* | Upregulation of *walKR* in vancomycin-resistant strains and inactivation of *walKR* cause increased sensitivity to vancomycin. | (*75-77*) |
| | T101M mutation in *walR* results in defective cell wall recycling and resistance to antibiotics targeting lipidII cycle. | (*78*) |
| | Co-occurrence of *walK* mutation with *rpoB* and *rpoC* mutants and resultant dual heteroresistance to daptomycin and vancomycin | (*48, 79-82*) |
| *Enterococcus faecalis* | *walK, rpoB* and *rpoC* mutations were found to be associated with in vivo daptomycin resistance | (*50, 83*) |
| *Corynebacterium glutamicum* | Deletion of *mtrA* and *mtrB* genes resulted in changed cell morphology and antibiotic susceptibilities | (*84*) |

**Table S3: Synergy of RIF with different drugs.**

| S. No. | Drug combination | Drug Class | Most synergistic area score | | |
|---|---|---|---|---|---|
| | | | ***M. tuberculosis*** | ***S. aureus*** | ***B. subtilis*** |
| 1. | Cefadroxil-RIF | Cell wall | 28.9 | 12.8 | 20.9 |
| 2. | Cephradine-RIF | Cell wall | 42.7 | NA | NA |
| 3. | Cefuroxime-RIF | Cell wall | NA | 12.4 | 19.7 |
| 4. | Meropenem-RIF | Cell wall | 19.4 | 5.2 | 10.6 |
| 5. | Faropenem-RIF | Cell wall | 15.5 | 15.5 | 14.0 |
| 6. | D-cycloserine-RIF | Cell wall | 6.2 | NA | NA |
| 7. | Ethambutol-RIF | Cell wall | 35.2 | NA | NA |
| 8. | Vancomycin-RIF | Cell wall | 1.8 | 23.7 | 4.3 |
| 9. | Teicoplanin-RIF | Cell wall | NA | 33.6 | 18.4 |
| 10. | Tunicamycin-RIF | Cell wall | NA | 28.2 | 6.4 |
| 11. | Isoniazid-RIF | Cell wall | 6.0 | NA | NA |
| 12. | MSO-RIF | Glutamine Biosynthesis | 1.7 | NA | NA |
| 13. | Moxifloxacin-RIF | DNA replication | 1.8 | 27.2 | 1.5 |
| 14. | Ciprofloxacin-RIF | DNA replication | 2.2 | 22.3 | 2.1 |
| 15. | Streptomycin-RIF | Translation | 17.3 | 2.2 | 5.3 |
| 16. | Fosfomycin-RIF | Cell wall | NA | 15.0 | 26.8 |

***Scale for synergistic area score: Less than -10: antagonistic, From -10 to 10: additive, Larger than 10: synergistic***

NA: not detected

**Table S4: Bacterial strains used in this work**

| S.No. | Bacterial Strains | Designation | Type | Reference |
|---|---|---|---|---|
| 1. | *Mycobacterium tuberculosis, H37Rv* | H37Rv | Wild type, lab strain | This work |
| 2. | *Mycobacterium tuberculosis, Erdman* | Erdman or rifS | Wild type, lab strain | (*85*) |
| 3. | *Mycobacterium tuberculosis, Erdman, S450L* | S450L or rifS | RIF resistant | (*85*) |
| 4. | *Mycobacterium tuberculosis, Erdman, H445Y* | H445Y or rifS | RIF resistant | (*85*) |
| 5. | *Mycobacterium tuberculosis, CDC155* | CDC155 | Wild type, lab strain | This work |
| 6. | *Mycobacterium tuberculosis, HN878* | HN878 | Wild type, lab strain | This work |
| 7. | *Mycobacterium tuberculosis, SA161* | SA161 | Wild type, lab strain | This work |
| 8. | *Mycobacterium tuberculosis, 17874* | 17874 | Wild type, clinical strain | (*61*) |
| 9. | *Mycobacterium tuberculosis, 20142* | 20142 | Wild type, clinical strain | This work |
| 10. | *Mycobacterium tuberculosis, 20389* | 20389 | Wild type, clinical strain | This work |
| 11. | *Mycobacterium smegmatis, mc*$^{2}$*155* | *M. smeg* | Wild type, lab strain | (*26*) |
| 12. | *Mycobacterium smegmatis, mc*$^{2}$*155; ftsZ::mCherry* | *ftsZ::* mCherry2B | Fluorescently labelled strain | (*25*) |
| 13. | *Mycobacterium smegmatis, mc*$^{2}$*155;rpoB::msfGFP* | *rpoB::*msfGFP | Fluorescently labelled strain | (*26*) |
| 14. | *Mycobacterium tuberculosis, ftsZ* knockdown | ftsZ KD | Mtb *ftsZ* KD | (*86*) |
| 15. | *Mycobacterium tuberculosis, mtrA* knockdown | *mtrA* KD | Mtb *mtrA* KD | (*9*) |
| 16. | *Mycobacterium tuberculosis, mtrA* complemented strain | *mtrA* Compli | Mtb *ftsZ* complementation strain | (*9*) |
| 17. | *Bacillus subtilis* subsp. 168 | *B. Sub or BS* | Wild type, lab strain | (*43*) |
| 18. | *Bacillus subtilis* subsp. 168; S487L | S487L | RIF resistant | (*43*) |
| 19. | *Bacillus subtilis* subsp. 168, H482Y | H482Y | RIF resistant | (*43*) |
| 20. | *Staphylococcus aureus*, N315 | SA | Wild type, lab strain | (*87*) |

**Table S5: sgRNAs used in this work**

| Figure Used | Targeted gene | Gene name | sgRNA targeting sequence (5'–3') | PAM (5'–3') |
|---|---|---|---|---|
| 1, 4, S4 | *rv0667* | *rpoB* | AGAACGACAACGACATCGACCC | GGAGAAG |
| 1, 4, S4 | *rv0668* | *rpoC* | GCAAGACCGATGCGGAGTTCATC | GAAGAAG |
| 3, S11 | *rv3246c* | *mtrA* | GGTGAGCATCACGATCGGAACA | CCGGAAT |
| 3, S12 | *rv2150c* | *ftsZ* | GCGGTGACAAACACCATG | TCGGCAC |
| | | | AGGGTTGCGCCGTTGGTGTGCA | ACGGCAC |

### Table S6: qPCR primers used in this work

| Figure Used | Target gene | Gene name | Forward primer (5'-3') | Reverse primer (5'-3') |
|---|---|---|---|---|
| 2, S10, S11 | *rv3246c* | *mtrA* | TGA TGC TCA CCG CAA AGA | CAG CGG TGT CAA CGA GAT |
| 2, S10, S11 | *rv3245c* | *mtrB* | CGA TTC GAG ACG TTG CTC A | ATC CAC CAG CAA CTC GAT AC |
| 2, S11, | *rv1477* | *ripA* | CCG GAT TCC GCG AGT TTA T | CCC AGC AAA CGA GTA CAA CA |
| 2, S10 | *rv1884c* | *rpfC* | GCA TGA CAA GAA TCG CCA AG | CTG CAG TCC GCC GTA TTT |
| 2, S10 | *rv0129c* | *fbpC* | GAG ACC TTC CTT ACC AGA GAG A | GTT GAG GAA GCC CGA CAA |
| 2, S10 | *rv0001* | *oriC (dnaA)* | CGG GAA TGC GGG TCA AAT A | GGT GTT GAA GGT GTG GAA GA |
| 2, S10 | *rv2163c* | *pbpB (ftsI)* | GTG TTC GGA AAG TCC TCC AA | ACC TTG GCC AAT AGG AAG ATT AG |
| 2, S10 | *rv2145c* | *wag31* | GCT TAC AAA CAC CGC CAA AG | GTA AGG CAT CGG CCT TCT C |
| 2, S10 | *rv3810* | *pirG* | TCC TCG GTG ATC CAA CAC T | TGC ATG ATC GAC GGC ATT AG |
| 2, S10 | *rv1221* | *sigE* | TAT CAC GAC CAT CAC GAC CT | TGC CGG AGA GCC GAT AA |
| 2, S10 | *rv2703* | *sigA* | CTA CGC TAC GTG GTG GAT TC | TGG ATT TCC AGC ACC TTC TC |
| 2, S10 | *rv0667* | *rpoB* | TGC TGC GTG CCA TCT T | GTC GGA GAT CTT GCG TTT CT |
| 2, S10 | *rv0668* | *rpoC* | GGG TTA TCC GTT CGT CAA CA | TGG TCG AGG ATC TCC TTC TT |
| 2, S10 | *rv1315* | *murA* | GCG GAG ATT CAG TTG GAG TT | CAC CGG TGA TGG TCA TTG T |
| 2, S10 | *rv2153c* | *murG* | GTG GCG GTG CCC TAT TT | ACC GGC AAC GCA TTC A |
| 2, S10 | *rv1018c* | *glmU* | TGG ATG ATC CCT TCG GCT A | CTC CTG TTG GGC GTT GTT |
| 2, S10 | *rv2145c* | *wag31* | TCA ACG AGC TGG ATC AAG AG | CTT TGG CGG TGT TTG TAA GC |
| 2, S10 | *rv2147c* | *sepF* | ACT ATC CAC CAC CGG GAT | GTG ATC TTC GAG AGC GGA TG |
| 2, S10 | *rv0050* | *ponA1* | AGA AGG CGG TTG CGA AAT A | CCA CCA GAG CAA ACA CCT TA |
| 2, S10 | *rv0017c* | *rodA* | TCG TTT CTG GTG GTG GTT TA | AAA GCG ACT GCA CGA TCT |
| 2, S10 | *rv1208* | *gpgS* | TGG CCT GGT CGA TGA ATT G | TCG ATG AAC ACC ACG ATG TC |
| 2, S10 | *rv3068c* | *pgmA* | TCA AGT ACA ACC CAC CCA AC | GGC AGG TCA TCG ACA TAG TG |
| 2, S10 | *rv2457c* | *clpX* | CGG GCT GGA GAA GAT CAT TTA | GAC TCT TTG TCC AGG TTG GT |
| 2, S10 | *rv3915* | *cwlM* | TTG CGC TCC TTG TAC TTT CT | TGC CAA GTC CCA CAA CAA |
| 2, S10 | *rv2220* | *glnA1* | GTT CCA GTC GAT CCA CGA AT | CAG TGC TGA TCA GGT AGT TCT C |

| | | | | |
|---|---|---|---|---|
| 2, S10 | *rv0993* | *galU* | GTTCTAGACCGTGCCATCTT | GTC ACG ATC CAA TGC AAA GTC |
| 2, S12 | *rv2150c* | *ftsZ* | CAA CTA CCT GGC CGT CAT C | AAG CAT CCA TCA GCG ATA CC |
| 2, S10, S11, s12 | *16s rRNA* | *16s rRNA* | GCC GTA AAC GGT GGG TAC TA | TGC ATG TCA AAC CCA GGT AA |
| 4 | *B.subtilis_walR* | *walR* | TATCGGCTCTCTCGTCATCTT | GGGTCATCACTTGTCCGATATG |
| 4 | *B.subtilis_walK* | *walK* | GACTGAAGGAGAACGCAGAAA | GCCGGGCTGTTTAAGAGAATA |
| 4 | *B.subtilis_groeL* | *BS_groEL* | CTTCTGATCGCTGAGGATGTT | CACCGAAACCAGGAGCTTTA |
| 4 | *S.aureus_walR* | *walR* | CACGTGTGAAAGCGAACTTAC | ATGCGTCTGGATAAATCACAATATC |
| 4 | *S.aureus_walK* | *walK* | GAAGCAGTCTAACCGTAGTCTAATC | GTCCTTACCACCGCCATAATC |
| 4 | *S.aureus_groeL* | *SA_groEL* | CAGCATTGATGCACGTGTAAG | TGCAACACCACCTGCTAAT |

**Table S7: Antibiotics used in this work**

| Antibiotic | Abbreviation | Product number/ Source |
|---|---|---|
| Rifampin | RIF | R3501 (Sigma Aldrich) |
| Fidaxomicin | FDX | SML1750 (Sigma Aldrich) |
| Myxopyronin B | MYXO | (*88*) |
| Bedaquiline (TMC-207) | BDQ | A12327 (Adooq Bioscience) |
| Cefadroxil | CD | C7020 (Sigma Aldrich) |
| Cephradine | CPH | C8395 (Sigma Aldrich) |
| Cefuroxime | CEF | C4417 (Sigma Aldrich) |
| Meropenem | MERO | 1392454 (Sigma Aldrich) |
| Faropenem | FARO | 122547-49-3 (Sigma Aldrich) |
| D-cycloserine | DC | 68-41-7 (Sigma Aldrich) |
| Ethambutol | ETM | J60695-06 (Alfa Aesar) |
| Vancomycin | VAN | 123409-00-7 (Alfa Chemistry) |
| Teicoplanin | TEIC | T0578 (Sigma Aldrich) |
| Tunicamycin | TUNI | T7765 (Sigma Aldrich) |
| Isoniazid | INH | 54-85-3 (Santa Cruz Biotechnology) |
| Methionine sulfoximine | MSO | M5379 (Sigma Aldrich) |
| Moxifloxacin | MOX | J66626 (Alfa Aesar) |
| Ciprofloxacin | CIPRO | 61-277-RF (Corning ) |
| Streptomycin | STREP | 3810-74-0 (Sigma Aldrich) |
| Fosfomycin | FOS | TCF0889 (TCI) |
